# Coopting T cell proximal signaling molecules enables Boolean logic-gated CAR T cell control

**DOI:** 10.1101/2022.06.17.496457

**Authors:** Aidan M. Tousley, Maria Caterina Rotiroti, Louai Labanieh, Lea Wenting Rysavy, Skyler P. Rietberg, Eva L. de la Serna, Guillermo Nicolas Dalton, Dorota Klysz, Evan W. Weber, Won-Ju Kim, Peng Xu, Elena Sotillo, Alexander R. Dunn, Crystal L. Mackall, Robbie G. Majzner

## Abstract

While CAR T cells have altered the treatment landscape for B cell malignancies, the risk of on-target, off-tumor toxicity has hampered their development for solid tumors because most target antigens are shared with normal cells^1,2^. Researchers have attempted to apply Boolean logic gating to CAR T cells to prevent on-target, off-tumor toxicity^3–7^; however, a truly safe and effective logic-gated CAR has remained elusive^8^. Here, we describe a novel approach to CAR engineering in which we replace traditional ITAM-containing CD3ζ domains with intracellular proximal T cell signaling molecules. We demonstrate that certain proximal signaling CARs, such as a ZAP-70 CAR, can activate T cells and eradicate tumors *in vivo* while bypassing upstream signaling proteins such as CD3ζ. The primary role of ZAP-70 is to phosphorylate LAT and SLP-76, which form a scaffold for the propagation of T cell signaling. We leveraged the cooperative role of LAT and SLP-76 to engineer <u>L</u>ogic-gated <u>I</u>ntracellular <u>N</u>etwor<u>K</u> (LINK) CAR, a rapid and reversible Boolean-logic AND-gated CAR T cell platform that outperforms other systems in both efficacy and the prevention of on-target, off-tumor toxicity. LINK CAR will dramatically expand the number and types of molecules that can be targeted with CAR T cells, enabling the deployment of these powerful therapeutics for solid tumors and diverse diseases such as autoimmunity^9^ and fibrosis^10^. In addition, this work demonstrates that the internal signaling machinery of cells can be repurposed into surface receptors, a finding that could have broad implications for new avenues of cellular engineering.

## Main Text

Chimeric antigen receptor (CAR) T cells have revolutionized the treatment of B cell malignancies, but are yet to make significant progress in treating solid tumors^1,2^. Lineage-derived B cell restricted antigens such as CD19 can be safely targeted with CAR T cells because depletion of B cells is not life threatening. This is not the case with solid tumors, where most overexpressed surface targets are also present on vital, normal tissues, creating the potential for on-target, off-tumor toxicity^11^. Thus, there is a dearth of antigens that can be safely targeted with CAR T cells for solid tumors^12^. As CARs are engineered to become even more potent and effective at recognizing low levels of antigen, the likelihood of on-target, off-tumor toxicity will further increase^13,14^. Thus, methods to apply Boolean logic to CAR T cells, endowing them with the ability to discriminate between normal and cancerous tissues, are essential to successfully target a large number of solid tumors.

Despite an intense focus on engineering more effective receptors^15–18^, CARs that are utilized today are very similar to the first iterations generated thirty years ago^19^. Almost all CARs contain a CD3ζ endodomain, the master switch (so-called ‘Signal 1’) for initiating the T cell signaling cascade^20,21^. Iterations and improvements have focused on the addition of ‘Signal 2’ (costimulatory domains)^22^ and ‘Signal 3’ (cytokine receptors)^23^, or on endowing cells with an ability to resist a suppressive tumor microenvironment^24^ or T cell exhaustion^25^. Other than manipulating the number of immunoreceptor tyrosine-based activation motifs (ITAMs) contained in the CD3ζ chain^13,26–28^, few attempts have been made to alter Signal 1 in CAR constructs.

The reliance on CD3ζ (or other ITAM-containing molecules) in CAR constructs has hampered the ability to apply Boolean logic gating to CAR T cells because ligation of the CAR alone triggers T cell activation. One method employed to overcome this limitation has been splitting the CD3ζ and costimulatory domains into CARs with different specificities so that maximal activity is only achieved when both targets are ligated (SPLIT CAR)^3–5^. However, CD3ζ-only constructs are capable of killing cells and have mediated on-target, off-tumor toxicity in clinical trials^29,30^. A more recently engineered system, SynNotch, utilizes a transcriptional circuit in which recognition of a first antigen drives expression of a traditional CD3ζ based CAR with specificity for a second target antigen^6^. While elegantly designed, this system does not escape the potential for on-target, off-tumor toxicity of bystander normal tissue once the CD3ζ CAR is expressed because the gene circuit is not immediately reversed^8^.

Here, we asked whether CARs utilize the same basic cellular signaling circuitry as the native T cell receptor (TCR). While we found that most proximal signaling molecules are necessary for CAR T cell activity, we also demonstrate that some molecules, such as ZAP-70 and PLCγ1, are sufficient themselves to initiate CAR T cell signaling, bypassing the need for CD3ζ. Armed with this finding, we drew on an understanding of T cell signaling networks to engineer the first true Boolean logic AND-gated CAR T cell through the pairing of LAT and SLP-76. This is the first system capable of restricting CAR T cell activity to encounter of two antigens in a direct, instantaneous, and reversible manner.

## Results

### CAR activity is dependent on the TCR machinery

Although all clinically validated CAR constructs contain CD3ζ, the master switch of activity for the T cell receptor (TCR)^20^, it is unknown whether CARs depend on the same proximal signaling networks as have been defined for the native TCR (Extended Data Figure 1a)^31^. We used CRISPR-Cas9 to individually knockout five proximal signaling molecules (Lck, Fyn, ZAP-70, LAT, and SLP-76) in primary human T cells expressing the CD19 CAR contained in tisagenlecleucel (Figure 1a, Extended Data Figure 1b) and measured degranulation (CD107a) and cytokine production (IL-2, TNF-α, and IFNγ) in response to antigen encounter (Figure 1b-e, Extended Data Figure 1c-e). While Fyn was expendable for CAR T cell function, in line with its overlapping role with Lck^32^ and dispensability for T cell development^33,34^, knockout of Lck, ZAP-70, LAT, or SLP-76 resulted in a near total ablation of CAR activity (Figure 1b-e, Extended Data Figure 1c-e). Taken together, these data indicate that CARs largely rely on the same proximal signaling networks that have been previously described for the native TCR.

**Figure 1:**
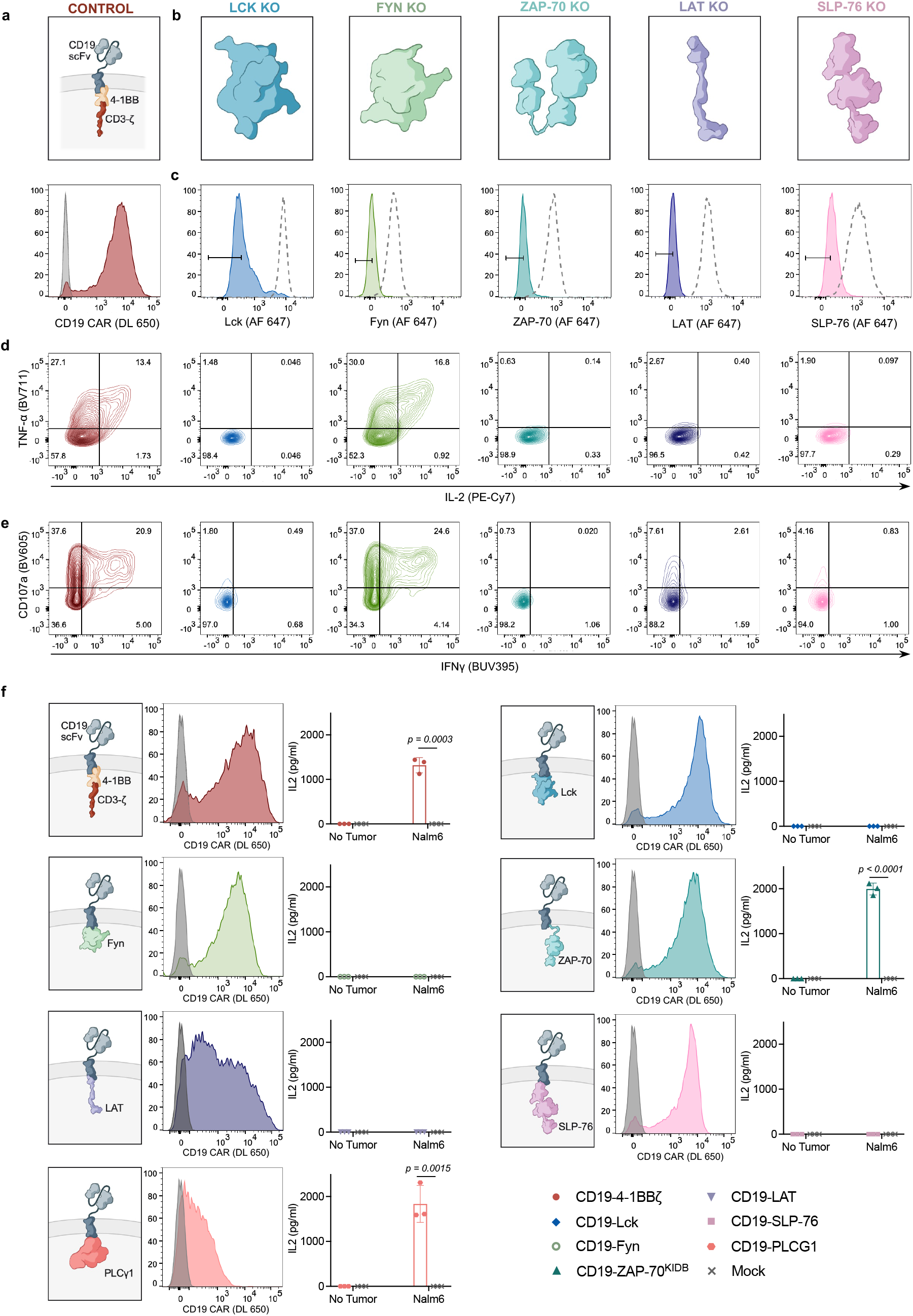
Proximal signaling molecules are necessary and sufficient to propagate CAR T cell activation. **a**, Schematic (top) and flow cytometric CAR expression (bottom) for unedited CD19-4-1BBζ CAR T cells. **b**, Schematics illustrating the five proximal signaling molecules targeted for CRISPR/Cas9-mediated knockout in CD19-4-1BBζ CAR T cells. **c**, Flow cytometric plots demonstrating knockout efficiencies for proximal signaling molecules illustrated in **b**. Dashed peaks represent the protein expression levels in unedited control cells. **d-e**, After CRISPR/Cas9-mediated knockout of proximal signaling molecules depicted in **b**, CD19-4-1BBζ CAR T cells were stimulated with Nalm6 tumor cells. Shown is flow cytometric data of TNF-α x IL-2 (**d**) and CD107a x IFNγ (**e**) in knockout populations designated in **c**. Data in **a-e** is representative of three independent experiments performed with different blood donors. **f**, Schematics (left), CAR expression (middle), and *in vitro* activity (right) of CD19-targeting CARs with proximal signaling endodomains. CARs incorporated full-length (Lck, Fyn, SLP-76, PLCγ1), intracellular (LAT), or truncated (ZAP-70, see **Extended Data Figure 1f**) domains. *In vitro* activity was assessed by measurement of IL-2 by ELISA in supernatant following co-culture of CAR T cells with CD19^+^ tumor cells (Nalm6). Data shown are mean values ± s.d of three experimental replicates. Representative of three independent experiments performed with different blood donors. *p* values were determined by the unpaired t-test (two-tailed).

### Downstream signaling proteins mediate effector functions as CARs

Having defined a set of proximal signaling molecules necessary for CAR T cell functionality, we next asked if any proximal signaling molecules themselves might be sufficient to induce T cell effector functions. We expressed six proximal signaling molecules (Lck, Fyn, ZAP-70, LAT, SLP-76, and PLCγ1) as CARs by tethering them to a transmembrane domain and a CD19 specific scFv (Figure 1f). Aside from LAT, these signaling molecules are natively located in the cytoplasm and do not contain a transmembrane domain, but expressed as transmembrane receptors when integrated into CARs (Figure 1f). Expression of a ZAP-70 kinase domain CAR required inclusion of interdomain B (IDB)^35^, a linker contained in the native molecule (ZAP-70KIDB, Extended Data Figure 1f-g). CARs bearing endodomains derived from ZAP-70 and PLCγ1 generated robust IL-2 in response to antigen encounter, while those containing Lck, Fyn, LAT, or SLP-76 endodomains did not (Figure 1f). We observed similar findings with CARs recognizing HER2, although HER2-PLCγ1 CAR activity was limited due to poor expression (Extended Data Figure 1h-i). Thus, some proximal signaling molecules are sufficient to initiate and propagate T cell activity and can be redeployed in surface receptors.

### ZAP-70 CARs demonstrate robust in vivo activity

To explore the utility of proximal signaling molecule CARs, we generated ZAP-70 CARs recognizing CD19, HER2, GD2, and B7-H3. Interestingly, we found significantly reduced expression of canonical T cell exhaustion markers on ZAP-70 CARs when utilizing scFvs that drive antigen independent tonic signaling such as GD2 and B7-H3 (Figure 2a-b, Extended Data Figure 2a)^36,37^. Consistent with this result, GD2-ZAP-70 T cells exhibited decreased antigen-independent IFNγ production (Extended Data Figure 2b). Although *in vitro* tumor cell killing and IL-2 production did not meaningfully differ between ZAP-70- and CD3ζ-based GD2 or B7-H3 CARs (Extended Data Figure 2c-d), B7-H3-ZAP-70 CARs eradicated tumors in a xenograft model of metastatic neuroblastoma in which B7-H3-4-1BBζ CARs were capable of only transient tumor control (Figure 2c-d). This enhanced anti-tumor activity was accompanied by substantially increased ZAP-70 CAR T cell expansion and persistence (Figure 2e, Extended Data Figure 2e). These data demonstrate that proximal signaling based CARs are capable of mediating robust antitumor activity *in vivo* and may be advantageous when utilized with scFvs demonstrating a high degree of tonic signaling.

**Figure 2:**
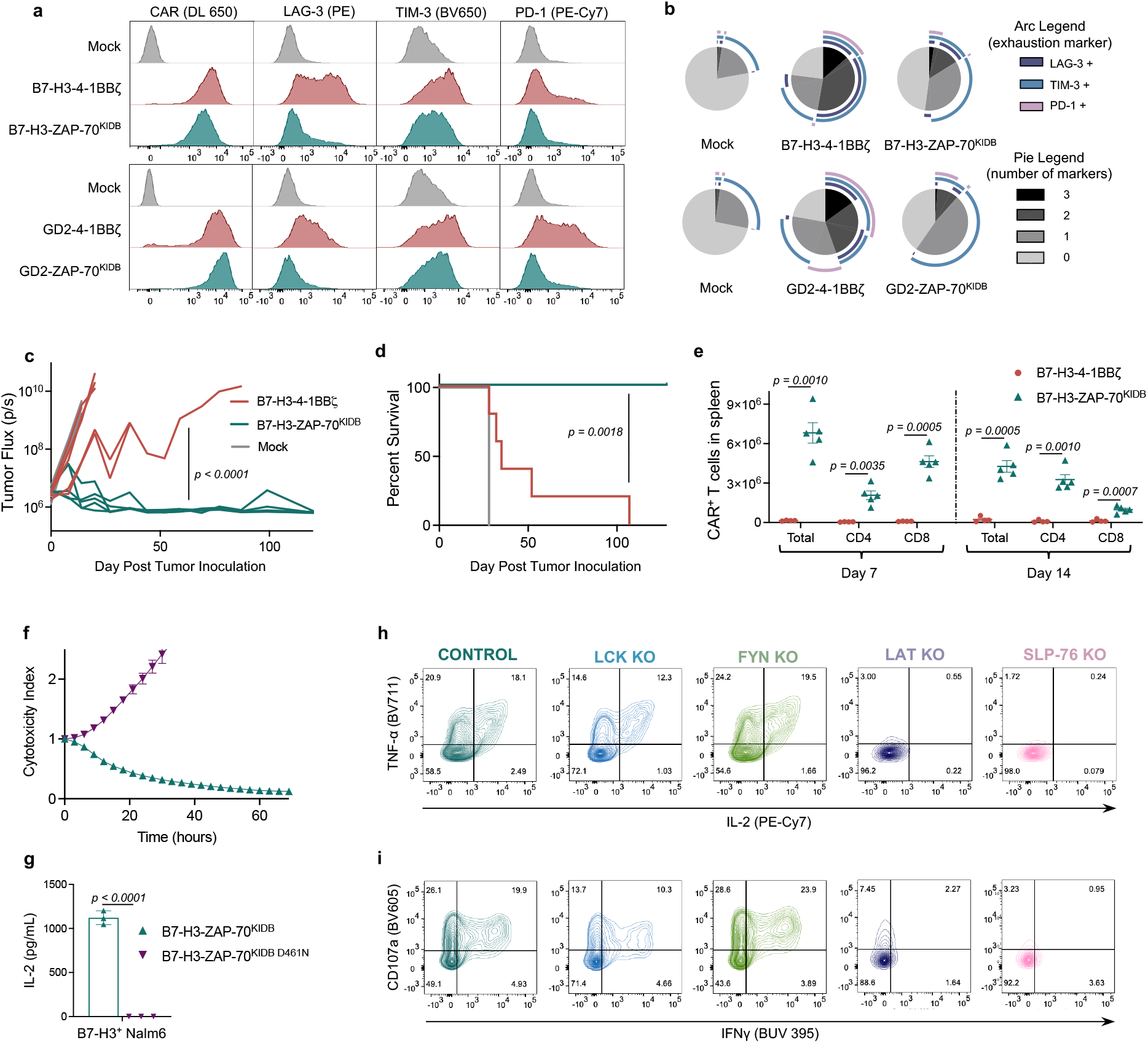
ZAP-70 CARs bypass upstream signaling elements and can exhibit enhanced antitumor activity. *a,* Representative flow cytometric plots of CAR, LAG-3, TIM-3, and PD-1 expression for T cells bearing B7-H3 or GD2-specific CARs containing 4-1BBζ or ZAP-70^KIDB^ endodomains (day 10 after T cell activation). **b**, Quantified SPICE (“Simplified Presentation of Incredibly Complex Evaluations”) plots from data show in **(a)**. Data in **a-b** is representative of four independent experiments with different T cell donors. **c-d**, NSG mice bearing CHLA-255-luciferase xenografts were treated intravenously with B7-H3-4-1BBζ or B7-H3-ZAP-70^KIDB^ CAR T cells. **(c)** Quantification of tumor progression for each individual mouse as measured by flux values acquired via bioluminescence imaging (BLI). (**d**) Survival curves for mice bearing tumors shown in **c.** Data is representative of three independent experiments with three different donors (n=5 mice per group in each experiment). *p* value in **c** was determined by repeated measures one way ANOVA with correction for multiple comparisons and **d** was determined by the Log-rank test. **e**, Absolute number of CAR T cells recovered from the spleens of CHLA-255-bearing mice on days 7 and 14 after treatment with B7-H3-4-1BBζ or B7-H3-ZAP-70^KIDB^ CAR T cells. Shown are mean values ± s.e.m. for n=4-5 mice per group per timepoint. Experiment was performed once at two timepoints. *p* values were determined by unpaired t-test (two-tailed) with Welch’s correction. **f**, Tumor cell killing of B7-H3^+^Nalm6-GFP cells co-cultured with B7-H3-ZAP-70^KIDB^ (± D461N mutation) CAR T cells at a 1:1 ratio of T cells to tumor cells. Shown are mean values ± s.d. of three experimental replicates. Representative of two independent experiments with different T cell donors. **g**, IL-2 secretion (as measured by ELISA) by B7-H3-ZAP-70^KIDB^ (± D461N mutation) CAR T cells following co-culture with B7-H3^+^ Nalm6 cells. Shown are mean values ± s.d. of three experimental replicates. *p* values were determined by the unpaired t-test (two-tailed). Representative of two independent experiments with different T cell donors. **h-i**, After CRISPR/Cas9-mediated knockout of the indicated proximal signaling molecules, B7-H3-ZAP-70^KIDB^ CAR T cells were stimulated with B7-H3^+^Nalm6 tumor cells. Shown is flow cytometric data of TNF-α x IL-2 (**h**) and CD107a x IFNγ (**i**) in knockout populations designated in **Extended Data Figure 3e**. Data in **h-i** is representative of three independent experiments performed with two different blood donors.

### ZAP-70 CARs depend on intrinsic kinase activity, bypassing CD3ζ and Lck

We then asked whether the activity of the ZAP-70 CAR is dependent on CD3ζ or other ITAMs found in the native TCR. CRISPR-Cas9 mediated knockout of the *TRAC* locus in ZAP-70 CAR T cells did not reduce tumor cell killing or IL-2 production (Extended Data Figure 3a-b), indicating that TCR ITAM domains are not required to initiate CAR T cell signaling. To understand the mechanism by which the ZAP-70 CAR propagates T cell signaling, we introduced a mutation that interrupts its kinase activity (D461N)^38^ and found that this abrogated both tumor cell killing and IL-2 production (Figure 2f-g, Extended Data Figure 3c). Furthermore, knockout of the downstream targets of ZAP-70, LAT or SLP-76, also abrogated ZAP-70 CAR T cell activity, while knockout of upstream molecules Lck or Fyn did not (Figure 2h-i, Extended Data Figure 3d-h). Thus, ZAP-70 CARs rely on activity of their kinase domain on downstream molecules and, unlike CD3ζ based CARs, can activate independently of TCR ITAMs and Src family kinase members.

### Pairing LAT and SLP-76 CARs allows for Boolean-logic AND-gating

Endogenous ZAP-70 is known to phosphorylate LAT and SLP-76, which then form a scaffold for PLCγ1 and other signaling molecules to propagate downstream effector functions^39^. Given that both ZAP-70 and PLCγ1 are sufficient to promote CAR-T cell activation, and that the ZAP-70 CAR is dependent on the presence of LAT and SLP-76, we hypothesized that we could similarly initiate T cell activity by clustering LAT and SLP-76 CARs to form a synthetic scaffold (Figure 3a). While T cells expressing either a CD19-LAT CAR or a HER2-SLP-76 CAR did not produce IL-2 in response to antigen encounter, T cells co-transduced with both CARs robustly responded to dual antigen encounter (Figure 3b). This stood in contrast to all other combinations of other inactive proximal signaling CARs (Lck, Fyn, SLP-76, and LAT), none of which responded to dual antigen encounter (Extended Data Figure 4a-b).

**Figure 3:**
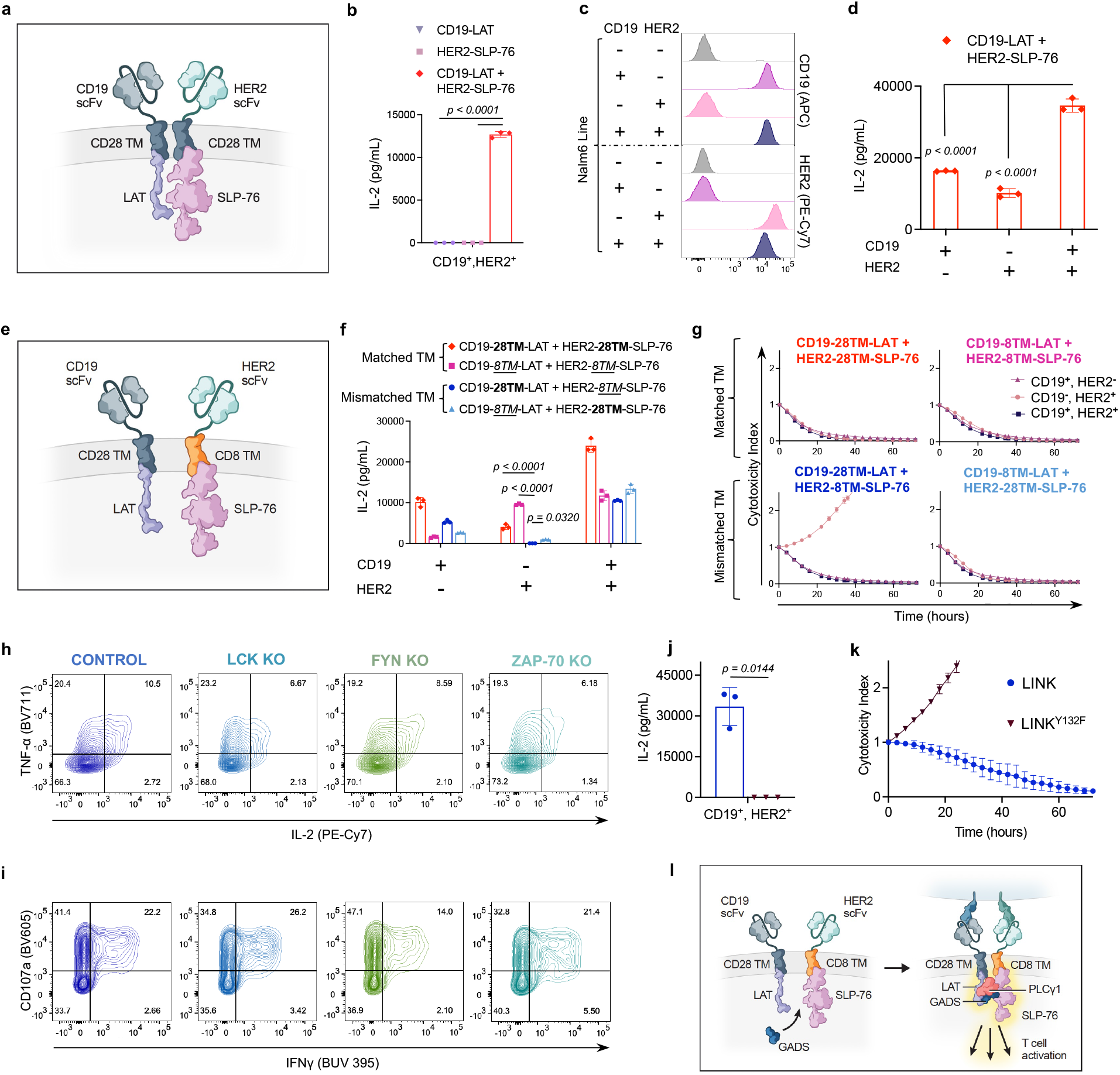
LAT and SLP-76 CARs bypass upstream signaling elements and function together as a Boolean logic AND-gate. **a,** Schematic illustrating LAT and SLP-76 CARs co-expressed on one T cell. **b,** IL-2 secretion (as measured by ELISA) by CD19-LAT, HER2-SLP-76, or CD19-LAT + HER2-SLP-76 CAR T cells following co-culture with HER2^+^ Nalm6 (CD19^+^, HER2^+^). Shown are mean values ± s.d. of three experimental replicates. Representative of eight independent experiments with different T cell donors. *p* values were determined by one way ANOVA with correction for multiple comparisons. **c,** Flow cytometry plots of CD19 and HER2 expression on engineered Nalm6 lines. **d,** IL-2 secretion (as measured by ELISA) by CD19-LAT + HER2-SLP-76 CAR T cells following co-culture with cell lines shown in **c**. Shown are mean values ± s.d. of three experimental replicates. Representative of eight independent experiments with different T cell donors. *p* values were determined by one way ANOVA with correction for multiple comparisons. **e,** Schematic illustrating LAT and SLP-76 CARs with mismatched hinge-transmembrane (TM) domains co-expressed on one T cell. **f,** IL-2 secretion (as measured by ELISA) by CD19-LAT + HER2-SLP-76 CAR T cells with the indicated hinge-transmembrane (TM) regions following co-culture with cell lines shown in **c**. Shown are mean values ± s.d. of three experimental replicates. Representative of three independent experiments with different T cell donors. *p* values were determined by one way ANOVA with correction for multiple comparisons. **g,** Tumor cell killing of cell lines shown in **c** co-cultured with CD19-LAT + HER2-SLP-76 CAR T cells with the indicated hinge-transmembrane (TM) regions at a 1:1 ratio of T cells to tumor cells. Shown are mean values ± s.d. of three experimental replicates. Representative of three independent experiments with different T cell donors. **h-i**, After CRISPR/Cas9-mediated knockout of the indicated proximal signaling molecules, LINK (CD19-28TM-LAT + HER2-8TM-SLP-76) CAR T cells were stimulated with HER2^+^Nalm6 (CD19^+^, HER2^+^) tumor cells. Shown is flow cytometric data of TNF-α x IL-2 (**h**) and CD107a x IFNγ (**i**) in knockout populations designated in **Extended Data Figure 5b**. Data in **h-i** is representative of two independent experiments performed with different blood donors. **j,** IL-2 secretion (as measured by ELISA) by CD19-28TM-LAT + HER2-8TM-SLP-76 (± LAT^Y132F^ mutation) CAR T cells following co-culture with HER2^+^Nalm6 (CD19^+^, HER2^+^) tumor cells. Shown are mean values ± s.d. of three experimental replicates. Representative of four independent experiments with different T cell donors. *p* value was determined by the unpaired t-test (two-tailed). **k,** Tumor cell killing of HER2^+^ Nalm6 (CD19^+^, HER2^+^) cells co-cultured with CD19-28TM-LAT + HER2-8TM-SLP-76 (± LAT^Y132F^ mutation) CAR T cells at a 1:1 ratio of T cells to tumor cells. Shown are mean values ± s.d. of three experimental replicates. Representative of four independent experiments with different T cell donors. **l**, Schematic illustrating the potential mechanism of LINK CAR activity.

Given their ability to respond to dual antigen expressing cells, paired LAT and SLP-76 CARs appeared to have the potential for Boolean logic AND-gating, in which CAR T cell activity is dependent on encounter of two distinct antigens, increasing their specificity in the clinic. However, while T cells co-transduced with both CD19-LAT and HER2-SLP-76 CARs responded to dual antigen encounter, they also demonstrated some degree of ‘leakiness,’ responding also to tumor cells expressing only one antigen (CD19 or HER2) (Figure 3c-d). Leakiness potentially increases the risk of CAR-mediated toxicity due to recognition of normal tissue expressing only one of the target antigens.

As both the LAT and SLP-76 component CARs contained a CD28 hinge/transmembrane (TM) domain, we hypothesized that these may homodimerize, bringing both CARs to the immune synapse even when only one antigen is engaged. Alternating the CAR TM domains between the constructs (CD28 and CD8, Figure 3e, Extended Data Figure 4c) reduced some of the single antigen activity, especially when T cells encountered HER2, the cognate antigen for the SLP-76 CAR (Figure 3f). The most promising combination (CD19-28TM-LAT + HER2-8TM-SLP-76) did not kill tumor cells expressing only HER2, but maintained some leaky activity, killing tumor cells expressing only CD19 (Figure 3g). We termed this combination **<u>L</u>**ogic-gated **<u>I</u>**ntracellular **<u>N</u>**etwor**<u>K</u>**(LINK) CAR and undertook further engineering to enhance its specificity. To reduce potential hetero- and homo-dimerization, we mutated the cysteine residues in the CD28 TM on the LAT CAR (2CA, Extended Data Figure 4d-e), which resulted in additional reduction of single antigen leakiness, but did not prevent killing of tumor cells expressing only CD19 (Extended Data Figure 4f-g).

### Targeted mutations establish AND-gate specificity

Given the persistent leakiness in the LINK platform, we undertook mechanistic studies to help guide our engineering approach. Knockout of ZAP-70, Lck, or Fyn did not abrogate T cell activity, demonstrating that LINK CARs bypass the upstream members of the proximal signaling cascade (Figure 3h-i, Extended Data Figure 5a-e). Furthermore, mutation of the tyrosine in LAT required for recruitment of PLCγ1 (Y132F)^40,41^ abrogated LINK CAR T cell activity (Figure 3j-k, Extended Data Figure 5f), demonstrating that downstream recruitment of PLCγ1 is essential for its function.

In native T cells, LAT and SLP-76 do not interact directly, but instead through adapter molecules from the Grb2 family such as GADS^41,42^. We hypothesized that upon dual antigen encounter, the LAT and SLP-76 CARs come together through association with GADS (or other Grb2 family members) to form a scaffold for PLCγ1 (Figure 3l), and that this association may have also been occurring in the absence of dual antigen ligation, causing leakiness. Therefore, we deleted the GADS binding sites in both the LAT (del171-233)^41^ and SLP-76 (del224-244)^43^ CARs (Figure 4a). When combined with the CD28 H/TM 2CA mutation detailed above (Extended Data Figure 6a-b), these targeted deletions resulted in a true AND-gate system, in which cytokine production (Figure 4b) and tumor cell killing (Figure 4c), as well as T cell degranulation and activation (Figure 4d, Extended Data Figure 6c) were largely eliminated except in response to dual antigen encounter. Targeted mutations to the tyrosine residues in LAT that interrupt its interaction with GADS (Y171F/Y191F)^41,44^ had a similar effect to the GADS truncations (Extended Data Figure 6d-e), further demonstrating the importance of the LAT/GADS interaction to effective engineering of the LINK CAR.

**Figure 4:**
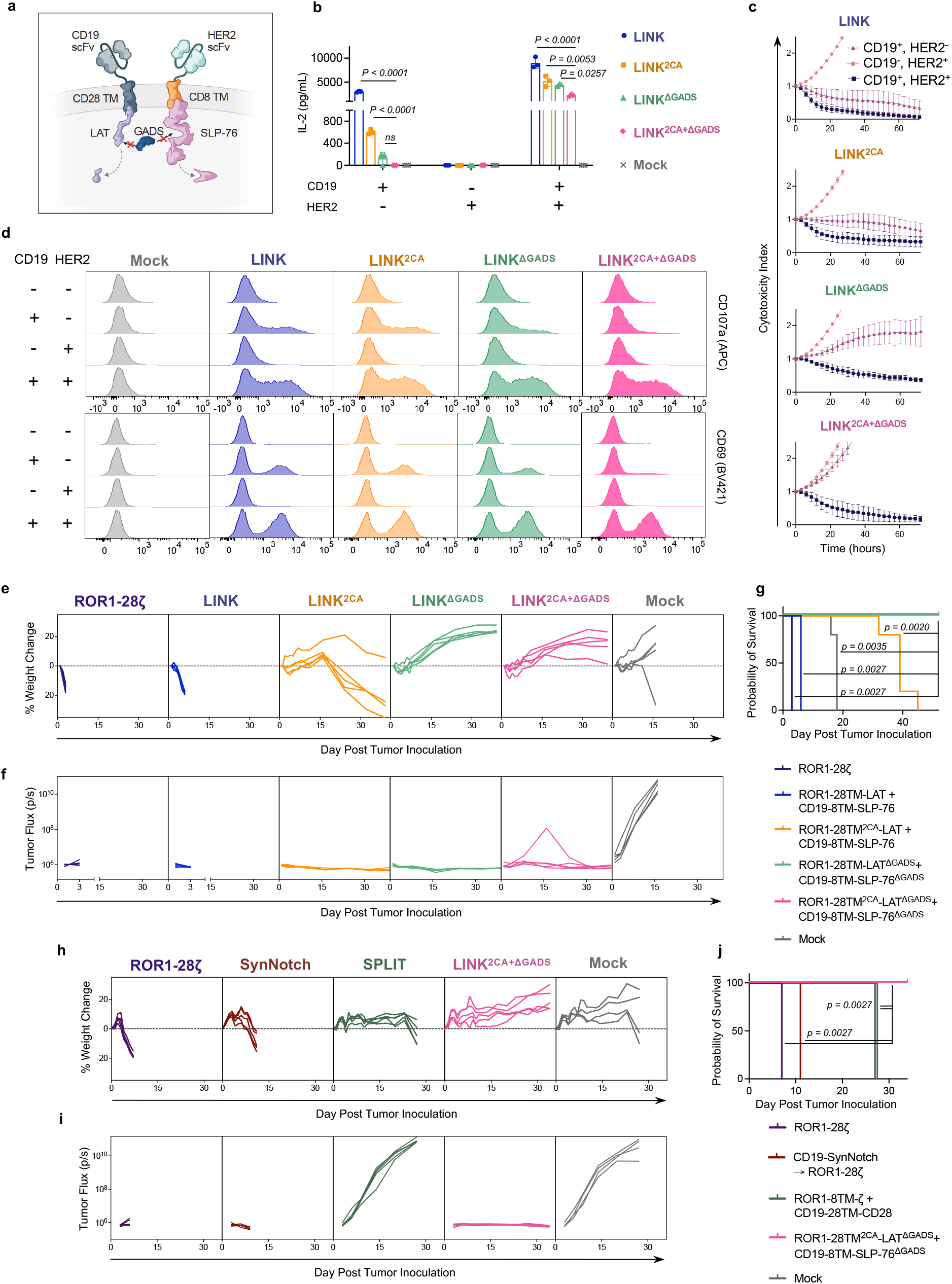
LINK CAR mediates tumor eradication while preventing on-target, off-tumor toxicity. **a,** Schematic illustrating truncation of the GADS-binding regions in the LAT and SLP-76 CAR endodomains. **b,** IL-2 secretion (as measured by ELISA) by indicated LINK CAR T cells following co-culture with cell lines shown in **Figure 3c**. Shown are mean values ± s.d. of three experimental replicates. Representative of five independent experiments with four different T cell donors. *p* values were determined by one way ANOVA with correction for multiple comparisons. **c,** Tumor cell killing of cell lines shown in **Figure 3c** by indicated LINK CAR T cells at a 1:1 ratio of T cells to tumor cells. Shown are mean values ± s.d. of three experimental replicates. Representative of five independent experiments with four different T cell donors. **d**, Flow cytometric plots of T cell degranulation (CD107a, top) and activation (CD69, bottom) on indicated LINK CAR T cells following co-culture with cell lines shown in **Figure 3c**. Representative of 5 independent experiments with four different T cell donors. **e-g**, NSG mice bearing ROR1^+^Nalm6-luciferase were treated with the indicated CAR T cells. **(e)** Weights for individual mice over time plotted as a percentage of the weight on day 0. **(f)** Quantification of tumor progression for each individual mouse as measured by flux values acquired via bioluminescence imaging (BLI). (**g**) Survival curves for mice bearing tumors shown in **f.** *p* values were determined by the Log-rank test. Data for **e-g** is representative of three independent experiments with two different blood donors each performed with n=5 mice per group. Note that in one of three experiments, the LINK^2CA^ mice did not lose enough weight to be euthanized. **h-j**, NSG mice bearing ROR1^+^Nalm6-luciferase were treated with the indicated CAR T cells and logic-gated CAR T cell systems. **(h)** Weights for individual mice over time plotted as a percentage of the weight on day 0. **(i)** Quantification of tumor progression for each individual mouse as measured by flux values acquired via bioluminescence imaging (BLI). **(j)** Survival curves for mice bearing tumors shown in **i.***p* values were determined by the Log-rank test. N=5 mice per group.

### LINK mediates tumor clearance and survival in a model on-target, off-tumor toxicity

To test the *in vivo* efficacy and specificity of the LINK CAR, we utilized a model of on-target, off-tumor toxicity mediated by a ROR1 specific CAR that recognizes both human and murine ROR1 (Extended Data Figure 7a-c)^8^ (*Labanieh et al., in revision*). On-target, off tumor recognition of ROR1 expressed on mouse lung tissues by CAR T cells causes toxicity manifested by rapid weight loss and death (Extended Data Figure 7a) *(Labanieh et al., in revision).*

Using isogenic cell lines (Extended Data Figure 7c), we confirmed the *in vitro* activity and dual antigen specificity of ROR1/CD19 LINK CARs (Extended Data Figure 7d-e), then proceeded to test multiple LINK CAR iterations *in vivo.* Mice bearing ROR1^+^, CD19^+^ Nalm6 xenografts were inoculated with ROR1 targeted CAR T cells. As we observed greater leakiness from the LAT CAR components in the early LINK CAR iterations, we first tested LINK CARs with specificity for ROR1 on LAT and CD19 on SLP-76. Similar to the conventional ROR1-CD28ζ CAR, the ROR1-CD28TM-LAT + CD19-CD8TM-SLP-76 CAR (LINK) mediated on-target toxicity, as evidenced by rapid weight loss and death. Mice that received ROR1-CD28TM^2CA^-LAT + CD19-CD8TM-SLP-76 CAR (LINK^2CA^) demonstrated longer survival, but toxicity eventually manifested after several weeks. However, LINK CARs in which the GADS interaction sites were deleted (either ROR1-CD28TM-LAT^ΔGADS^+ CD19-CD8TM-SLP-76^ΔGADS^ [LINK^ΔGADS^] or ROR1-CD28TM^2CA^-LAT^ΔGADS^+ CD19-CD8-SLP-76^ΔGADS^ [LINK^2CA+ΔGADS^]) mediated complete tumor cell clearance without any evidence of on-target, off tumor toxicity (Figure 4e-g). We also reversed the orientations of these CARs (now with ROR1 specificity on SLP-76 and CD19 on LAT) and found that even in this orientation, prevention of toxicity required use of the optimized LINK^2CA+ΔGADS^CAR (CD19-CD28TM^2CA^-LAT^ΔGADS^+ ROR1-CD8TM-SLP-76^ΔGADS^, Extended Data Figure 8a-e).

There have been several previous attempts in the literature to generate AND gate CARs. These include the ‘SPLIT’ CAR system, in which the CD3ζ and costimulatory domains are split into CARs with different specificities^3–5^, as well as gene-circuits such as the SynNotch system^6^. In the SynNotch system, response to encounter of a first antigen (antigen A), a synthetic Notch receptor releases a transcription factor that then drives expression of a traditional cytolytic CAR with specificity for a second antigen (antigen B). Thus, SynNotch cells are primed by antigen A to recognize and kills cells expressing antigen B. While elegantly designed to program transcriptional responses in T cells, this system is not a true AND-gate because cells that are primed by antigen A can then attack any tissue expressing antigen B (Extended Data Figure 9a). Previous work has demonstrated that SynNotch does not prevent on-target, off-tumor toxicity in mouse models^8^.

We compared our LINK CAR platform to both SynNotch (CD19-SynNotch à ROR1-CD28ζ) and SPLIT (ROR1-8TM-ζ+ CD19-28TM-CD28) CARs (Extended Data Figure 9b) in the toxicity model described above. We confirmed that the SynNotch system was appropriately primed by encounter of CD19 to express the ROR1 CAR (Extended Data Figure 9c) and that its specificity was largely restricted to dual antigen encounter *in vitro* (Extended Data Figure 9d). However, the SynNotch CAR mediated significant on-target, off-tumor toxicity similar to that which was observed with a traditional ROR1-CD28ζ CAR (albeit slightly delayed). The SPLIT CAR system mediated neither toxicity nor detectable tumor control, performing similarly to MOCK T cells, although it did demonstrate some *in vitro* activity (Extended Data Figure 9e). In contrast, LINK^2CA+ΔGADS^ CAR mediated complete tumor control without any signs of on-target, off-tumor toxicity (Figure 4h-j). Thus, the LINK CAR platform is capable of specific and effective antitumor activity while preventing fatal on-target, off-tumor toxicity.

## Discussion

CAR T cells have become an essential tool in the treatment of B cell malignancies and are a curative, life-saving therapy for many patients^2,45–53^. However, the deployment of CAR T cells to treat solid tumors has been slow and marked by failures^1,54^. One major obstacle is the dearth of tumor specific targets not shared with vital, normal tissues^11,12^. While CARs can sometimes demonstrate a therapeutic window between tumors expressing very high levels of antigen and normal cells expressing the same target at lower levels^13,37,55–58^, as more potent CAR T cells are engineered, more frequent on-target, off tumor toxicity is likely to emerge in clinical studies.

We have generated the first true Boolean-logic AND-gated CAR T cells by coopting the cell’s LAT/SLP-76 scaffold that is required for T cell signaling. By leveraging native T cell signaling biology, we were able to achieve a highly specific and portable system for restricting T cell responses to dual antigen encounter. Several other AND-gate CAR systems have been engineered, including SynNotch^6^ and SPLIT CAR^3–5^, but as demonstrated by others^8^ and in our data, these have shortcomings in both specificity and efficacy *in vivo*. Two additional AND-gate systems which require co-administration of foreign proteins to redirect CAR T cells based on antigen expression have been proposed and tested *in vitro* (LOCK-R^59^ and SUPRA CAR^60^). Small protein therapeutics are technically challenging to administer in the clinic due to their pharmacokinetics and may have limited ability to effectively traffic into tissues. Furthermore, completely synthetic proteins are potentially highly immunogenic. In contrast, the signaling components of the LINK CAR are fully human and the system can be readily engineered to any specificity in a modular fashion. Identification of ideal AND-gate targets is a growing field in oncology^61,62^, and this system is poised to alter the landscape of what molecules can be safely targeted with CAR T cells. Additional work utilizing patient samples and single cell technologies^62–64^ will be necessary to identify safe and effective targets for LINK CAR T cells.

We engineered LINK CARs after observing that intracellular, proximal signaling molecules such as ZAP-70 and PLCγ1 can function as signaling domains in synthetic transmembrane receptors. This work demonstrates that synthetic receptors can repurpose cytosolic kinases and other signaling machinery to control intracellular processes. This discovery may be broadly applicable across cell types and lead to the development of new cellular therapies apart from those using T cells.

Proximal signaling CARs, including ZAP-70 and LINK CARs, bypass the upstream components of the TCR machinery, but it remains undefined how these constructs are phosphorylated and whether the transcriptional programs they induce are similar to those induced by traditional CD3ζ containing CARs. These newly designed CARs also bypass a highly evolved system for kinetic proofreading that reduces signal transduction in response to low affinity interactions and prevents autoimmunity driven by self-reactive TCRs^65^. ZAP-70 CARs demonstrate some advantages in our models including reduced tonic signaling and T cell exhaustion as well as increased expansion. It is possible that some ZAP-70 CARs outperform CD3ζ CARs because their signal strength is better calibrated^28^ or because, by bypassing CD3ζ and Lck, ZAP-70 CARs may evade inhibitory ligands and other regulatory mechanisms in T cells^66^. Future work will define the basis for enhanced ZAP-70 CAR activity and assess whether the same advantages hold true for LINK and PLCγ1 CARs.

In summary, we have engineered a toolbox of novel CARs using proximal signaling molecules that demonstrate enhanced functionality and specificity, including a robust system for instantaneous and reversible Boolean logic AND-gating that outperforms previously published systems. By repurposing cytosolic molecules into CARs, we can control cellular activity in a highly functional but unanticipated manner. Our tools offer scientists and physicians newfound ability to control and enhance CAR T cell functionality in the clinic. These advances may have broad impact, not only in the field of cancer immunotherapy, but also as researchers extend CAR T cells to diseases such as autoimmunity and develop new classes of cellular therapies.

## Methods

### Construction of CAR constructs

CD19-4-1BBζ, HER2-4-1BBζ, B7-H3-4-1BBζ, and GD2-4-1BBζ CAR constructs were previously described^13,36,37^. CAR constructs were generated with a mix of restriction enzyme and In-Fusion HD Cloning (Takara Bio) using codon optimized gBlocks purchased from Integrated DNA Technologies. All CARs were cloned into MSGV1 vectors unless otherwise indicated. For Lck, Fyn, ZAP-70 and SLP-76 CARs, the full-length proteins were codon optimized and used. For LAT CARs, the intracellular sequence of LAT (residues 28-262) was similarly codon optimized and cloned. To generate CD19-ZAP-70^Kinase^ and CD19-ZAP-70^KIDB^ CARS, segments of ZAP-70 comprised of the Interdomain B and kinase domain regions (residues 255-600) or the kinase domain only (residues 338-600) were utilized. Mutations and deletions in these CARs were introduced via In-Fusion cloning on PCR fragments.

The ROR1 scFv was derived from a humanized clone F antibody (US Patent 20200405759A1), generated as a gBlock and cloned into CAR backbones. A VSV-g tag was added to the ROR1 scFv at the N-terminus after the signal sequence via In-Fusion Cloning to improve detection of ROR1-targeting CARs by flow cytometry. SPLIT CARs were generated by separating CD3ζ and CD28 domains into separate CARs using PCR and In-Fusion cloning.

The CD19 SynNotch construct was generated by PCR-amplifying the CD19 scFv, Notch extracellular domain, Notch transmembrane domain, Gal4, and VP64 from Addgene plasmid #79125 and cloning the product into the MSGV1 vector.

The SynNotch-inducible ROR1-28ζ construct was generated via insertion of a DNA sequence, from 5’ to 3’, encoding GAL4 UAS response elements from Addgene plasmid #79123, a minimal CMV promoter, GM-CSF leader sequence, VSVg tag, ROR1-CD28ζ CAR, woodchuck hepatitis virus post-transcriptional regulatory element (WPRE), EF1α promoter, and mTagBFP2 into a lentiviral backbone plasmid (System Biosciences # TR012PA).

All constructs were verified by DNA sequencing (Elim Biopharmaceuticals).

### Generation of cell lines

The Nalm6-GFP Luciferase B-ALL cell line was obtained from S. Grupp (University of Pennsylvania, Philadelphia, PA), CHLA-255 from R. Seeger (Keck School of Medicine, USC), and 143B from ATCC. The generation of B7-H3^+^, HER2^+^, GD2^+^, ROR1^+^ Nalm6 lines as well as CD19KO Nalm6 were previously described^25,37,67^ (*Labanieh et al., in revision).* All tumor cell lines were cultured in complete RPMI media supplemented with 10% FBS, 10 mM HEPES, 2 mM GlutaMAX, 100 U/mL penicillin, and 100 μg/mL streptomycin (Gibco).

To generate isogenic Nalm6-GL cell lines expressing CD19, HER2, and ROR1, Nalm6 cells were virally transduced with retroviral or lentiviral vectors encoding cDNA for the antigen of interest. Expression was verified via flow cytometry, and cells were FACS sorted on Stanford FACS Core Shared FACSAria cytometers (BD Biosciences). Cell lines were sorted to obtain matching target expression between lines.

### Production of retroviral and lentiviral supernatant

Retroviral supernatant was generated via transient transfection of 293GP cells, as previously described^36^. In brief, 6-7 x 10^6^ 293GP cells were added to 100 mm Poly-D-Lysine-coated plates in complete DMEM media supplemented with 10% FBS, 10 mM HEPES, 2 mM GlutaMAX, 100 U/mL penicillin, and 100 μg/mL streptomycin (Gibco). The following day, cells were cotransfected with 9 μg vector plasmid and 4.5 μg RD114 with Lipofectamine 2000 (Invitrogen) in Opti-MEM media (Gibco). The media was replaced after 24 hours and harvested at 48 and 72 hour time points. Viral supernatant was frozen at −80°C for long term storage.

Lentiviral supernatant was generated as previously described^67^. In brief, 6-7 x 10^6^ 293T cells were added to 100 mm Poly-D-Lysine-coated plates in complete DMEM media supplemented with 10%FBS, 10 mM HEPES, 2 mM GlutaMAX, 100 U/mL penicillin, and 100 μg/mL streptomycin (Gibco). The following day, cells were co-transfected with 9 μg vector plasmid, 9 μg pRSV-Rev, 9 μg pMDLg/pRRe, and 3.5 μg pMD2.G with Lipofectamine 2000 (Invitrogen) in Opti-MEM media (Gibco). The media was replaced after 24 hours and harvested at 48 and 72 hour time points. Viral supernatant was frozen at −80°C for long term storage.

### PBMC and T cell isolation

Healthy donor buffy coats, leukopaks or Leukocyte Reduction System (LRS) chambers were obtained through the Stanford Blood Center under an IRB-exempt protocol. Peripheral blood mononuclear cells were isolated using Ficoll-Paque Plus (GE Healthcare, 17-1440) density gradient centrifugation according to the manufacturer’s instructions and cryopreserved with CryoStor CS10 freeze media (Sigma-Aldrich) in 1-5 x 10^7^ cell aliquots. In some experiments, T cells were isolated using the RosetteSep Human T Cell Enrichment kit (Stem Cell Technologies) according to the manufacturer’s protocol.

### CAR T cell transduction and culture

CAR T cells were generated and cultured as previously described^67^. Briefly, cryopreserved PBMCs or T cells were thawed on day 0 and cultured with Human T-Activator αCD3/CD28 Dynabeads (Gibco) at 3:1 bead:cell ratio in AIM-V media (Gibco) supplemented with 5% FBS, 10 mM HEPES, 2 mM GlutaMAX, 100 U/mL penicillin, 100 μg/mL streptomycin, and 100 U/mL recombinant human IL-2 (Peprotech).

Retroviral/lentiviral transductions were performed on days 3 and 4 post activation on retronectin (Takara) coated non-tissue culture treated plates. Wells were coated with 1 mL of 25 μg/mL retronectin in PBS overnight, then blocked with 2% BSA in PBS for 15 minutes prior to transduction. 1 mL of thawed retroviral supernatant per CAR construct was added and plates were then centrifuged at 3,200 RPM at 32°C for 2-3 hours. Viral supernatant was discarded and 0.5 x 10^6^ T cells were added to each well in 1 mL of complete AIM-V media.

On day 5 post-activation, CD3/CD28 beads were removed with a magnet, and CAR T cells were maintained in culture with AIM-V media changes every 2-3 days at a density of 0.3 x 10^6^ cells/mL.

### Knockout of proximal signaling molecules

Proximal signaling molecule expression was disrupted with CRISPR-Cas9 mediated gene disruption. sgRNA for TRAC was previously described^13^. All other sgRNAs were designed using the Knockout Design Tool (Synthego). The sgRNA target sequences (5’ to 3’) used were: Lck-CTTCAAGAACCTGAGCCGCA Fyn-TGGAGGTCACACCGAAGCTG, AGAAGCAACAAAACTGACGG ZAP-70-CAGTGCCTGCGCTCGCTGGG, CCTGAAGCTGGCGGGCATGG LAT-CACACACAGTGCCATCAACA, CGTTTGAACTGGATGCCCCT SLP-76-AATAGTCAGCAAGGCTGTCG, GAAGAAGTACCACATCGATG TRAC-GAGAATCAAAATCGGTGAAT

Editing was performed on T cells after Dynabead removal five days after T cell activation. T cells were resuspended in P3 buffer (0.75-1 x 10^6^ cells per 18 μl P3) from the P3 Primary Cell 4D-Nucleofector X Kit S (Lonza). 10 μg/μl Alt-R S.p. Cas9 Nuclease V3 and Nuclease-free Duplex Buffer (IDT) were mixed in a 1:2 ratio; the ensuing mixture was then incubated at a 1:1 ratio with 120 pmol sgRNA for approximately 10 minutes. 18 μl of cells were mixed with 2 μl of sgRNA:Cas9 ribonucleoprotein complexes, then electroporated in 16-well cuvette strips under the E0-115 program. CAR T cells were quickly transferred into 200 μl of compete AIM-V media and allowed to incubate at 37°C for 24 hours before being moved to larger volumes and cultured as described above. For knockouts conducted using multiple sgRNAs, each sgRNA was incubated with Cas9/Duplex mix at a 1:1 ratio separately, then 2 μl of each mix was added to 18 μl of cells in P3 buffer (if two sgRNAs are used, the final volume was 22 μl). Knockout efficiency was assessed via intracellular flow cytometry upon expansion of the edited cells (as below).

### Flow cytometry

Cells were washed with FACS buffer (2% FBS in PBS) before staining. Staining was performed in FACS buffer for 20 minutes at 4°C. If a live/dead stain was used, 1X Fixable Viability Dye eFluor 780 (eBioscience) was also added to the staining mix. Cells were then washed once with FACS buffer on a BD Fortessa. FACSDiva software (BD) was used for Fortessa data collection.

T cells were assessed for CAR expression on the same day they were used for *in vitro/in vivo* assays. CD19-targeting CARs were detected using the anti-CD19 CAR idiotype antibody provided by L. Cooper (MD Anderson Cancer Center)^68^. GD2-targeting CARs were detected using the 1A7 anti-14G2a idiotype antibody obtained from National Cancer Institute. HER2, B7-H3, and human and mouse ROR1-targeting CARs were detected using human HER2-Fc, B7-H3-Fc, and human or mouse ROR1-Fc recombinant proteins (R&D). Idiotype antibodies and Fc-proteins were fluorophore-conjugated using DyLight 650 or DyLight 488 Microscale Antibody Labeling Kits (Invitrogen). ROR1 scFvs also contained a VSV-g tag that were detected by anti-VSV-g polyclonal antibody (FITC, Abcam).

The following antibodies were used for T cell staining:

CD69 (BV421, Clone FN50, BioLegend), CD107a (BV605 or APC, Clone H4A3, BioLegend), CD4 (BUV 737 or BUV 395, Clone SK3, BD Biosciences), CD8 (BUV 805, Clone SK1, BD Biosciences), CD3 (BUV 496, Clone UCHT1, BD Biosciences), CD45 (PerCP-Cy5.5, Clone HI30, Invitrogen), PD-1 (PE-Cy7, Clone EH12.2H7, BioLegend), TIM-3 (BV510 or BV650, Clone F38-2E2, BioLegend), LAG-3 (PE, Clone 3DS223H, Invitrogen), IgG1 κ isotype (PE, Clone 11711, R&D Systems), IgG1 κ isotype (PE-Cy7, Clone MOPC-21, BioLegend), IgG1 κisotype (BV650, Clone MOPC-21, BioLegend).

The following antibodies were used for tumor cell staining:

CD19 (APC, Clone HIB19, BioLegend), HER2 (PE-Cy7, Clone 24D2, BioLegend), and ROR1 (PE-Cy7, Clone 2A2, BioLegend).

SPICE (“Simplified Presentation of Incredibly Complex Evaluations”) plots were generated by calculating LAG-3/TIM-3/PD-1 CAR T cell populations on FlowJo and importing into SPICE software^69^.

Representative gating strategies for flow cytometric assays are shown in Extended Data Figure 10.

### Intracellular protein staining and intracellular cytokine assays

Staining was performed following the manufacturer’s protocol for the Foxp3/ Transcription Factor Staining Buffer Set (eBioscience). A fixable viability dye was added (eFluor 780, eBioscience), and relevant extracellular markers were stained prior to the fixation step. Following permeabilization, cells were stained with 0.1μg/100,000 cells of anti-Lck/Fyn/ZAP-70/LAT/SLP-76 antibodies. Isotype controls were utilized for gating.

For intracellular cytokine assays, CAR T cells (day 18 post activation) were incubated with target cells at a 1:1 ratio for 5 hours in the presence of 1X Monensin (eBioscience) and 0.75 μl anti-CD107a antibody/test. Following stimulation, the staining buffer set protocol was performed. 0.75μl/condition of anti-cytokine antibodies were added following permeabilization.

The following antibodies were used for intracellular staining:

Lck (Alexa Fluor 647, Clone Lck-01, BioLegend), Fyn (Alexa Fluor 647, Clone FYN-01, Novus Biologicals), ZAP-70 (Alexa Fluor 647, Clone A16043B, BioLegend), LAT (Alexa Fluor 647, Clone 661002, R&D Systems), SLP-76 (Alexa Fluor 647, Clone H3, BD Biosciences), IgG1 κ isotype (Alexa Fluor 647, Clone MOPC-21, BioLegend), IgG2a κ isotype (Alexa Fluor 647 Clone MOPC-173, BioLegend), IgG2b κ isotype (Alexa Fluor 647, Clone MPC-11, BioLegend), IL-2 (PE-Cy7, Clone MQ1-17H12, BioLegend), IFNγ (BUV 395, Clone B27, BD Biosciences), TNF-α (BV711, Clone MAb11, BioLegend).

### Cytotoxicity assays

CAR^+^ T cells (day 10 post activation) were co-cultured with 50,000 tumor cells at either 1:1 or 1:2 T cell to target cell ratios (E:T) in complete RPMI on 96-well flat bottom plates. Co-cultures were incubated at 37°C and imaged with an Incucyte S3 Live-Cell Analysis System (Sartorius) for approximately 72 hours. The basic analyzer feature on the Incucyte S3 software was used to quantify killing of GFP^+^ Tumor Cells by measuring the Total Green Object Integrated Intensity over time. Fluorescent values were normalized to the initial measurement at time 0.

### Cytokine assays

1 x 10^5^ CAR^+^ T cells (day 10 post activation) were co-cultured with tumor cells in a 1:1 E:T ratio in complete RPMI media and incubated at 37°C for approximately 24 hours. For cells stimulated with anti-CD3/anti-CD28 antibody coated DynaBeads (Gibco), beads were added to cells in a 3:1 ratio. After stimulation, the supernatants were collected, and IL-2 or IFNγ were measured by ELISA following the manufacturer’s protocol (BioLegend). Absorbances were measured with a Synergy H1 Hybrid Multi-Mode Reader (BioTek).

### T cell activation assays

For activation assays assessing CD69 and CD107a expression, 1 x 10^5^ double positive CAR T cells (day 10 post activation) were co-cultured with single CD19^+^, HER2^+^, and CD19^+^HER2^+^leukemia cells at 1:1 E:T ratios in complete RPMI media at 37°C for 4 hours in the presence 0.75 μl anti-CD107a antibody and 1X monensin (eBioscience). Cells were then washed with cold FACS buffer and staining was performed as described above.

### *In vivo* xenograft models

All animal studies were carried out according to Stanford Institutional Animal Care and Use Committee-approved protocols. Immunodeficient NOD-*scid* IL2Rg^null^ (NSG, NOD.Cg-PrkdcscidIl2rgtm1Wjl/SzJl) mice were purchased from The Jackson Laboratory or bred in house. 6-to-10-week-old mice were inoculated with 1 x 10^6^ CHLA-255 cells 7 days prior to T cell injection or 1 x 10^6^ ROR1 ^+^Nalm6-GL cells 1-3 days prior to T cell injection via intravenous injection (200 μl PBS per injection). Mice were randomized to ensure even tumor burden between experimental and control groups prior to treatment.

CAR T cells were injected intravenously; CHLA-255-bearing mice received 3 x 10^6^ (for efficacy/survival experiments) or 1 x 10^7^ (for proliferation/persistence experiments) CAR^+^ T cells (day 10 after activation), and ROR^+^Nalm6-bearing mice received 6-8 x 10^6^ double CAR+ T cells. Mice were monitored for disease progression 1-2 times per week via bioluminescence imaging (BLI) using an IVIS imaging system (Perkin Elmer). For CHLA-255 models, mice were humanely euthanized when they demonstrated morbidity or developed large solid tumor masses. For leukemia models, mice were humanely euthanized when they demonstrated morbidity or exhibited hind-leg paralysis. For the ROR1-CAR on-target/off-tumor toxicity model, mice were weighed frequently and humanely euthanized if their weight rapidly dropped 20%, or if they exhibited significant signs of distress (hunched posture, impaired mobility, rough coat, shivering).

### Assessment of *T cell* expansion *in vivo*

To assess CAR T cell expansion *in vivo*, mice were sacrificed at one- and two-week timepoints following T cell infusion. Spleens and bone marrow were harvested, counted, and stained for CD4, CD8, CD45, and B7-H3 CAR as described above. T cells were gated as GFP- and CD45+. The percentages of CAR^+^ cells and cell counts were used to calculate the absolute number of T cells present in the spleen.

### Statistical Analysis

Statistical analyses were performed with Excel version 16.53 (Microsoft) and GraphPad Prism 9.1.0 (GraphPad). Figure legends denote group mean values ± s.d. or s.e.m. Unless otherwise noted, analyses testing for significant differences between groups were conducted with unpaired two-tailed t-tests (when comparing two groups) or one-way ANOVA with multiple comparison correction (when comparing more than two groups). *In vivo* survival curves were compared with the log-rank Mantel-Cox test. *In vivo* tumor growth was compared with repeated-measures ANOVA with multiple comparisons. *In vivo* T cell proliferation was compared using unpaired two-tailed t-tests with Welch’s correction. *p* < 0.05 was considered statistically significant.

## Acknowledgements

This work was funded in part by the NIH Director’s New Innovator Award (DP2 CA272092 to R.G.M.) and the Parker Institute for Cancer Immunotherapy (R.G.M., C.L.M). E.W.W. is supported by a Bridge Fellow Award from the Parker Institute for Cancer Immunotherapy. The authors thank Drs. Howard Chang and Ansuman Satpathy for their review of the manuscript and insightful comments. The authors also thank SciStories (Sigrid Knemeyer, Vivian Yeung, and Su Min Suh) for providing the schematics (Figures 1, 3, and 4, and Extended Data Figures 1, 4, 6, 7, and 9) and consulting on figure design.

## Data Availability Statement

The datasets generated during this study will be uploaded with the final manuscript. CAR constructs will be made available through Material Transfer Agreements.

---

**Extended Data Figure 1:**
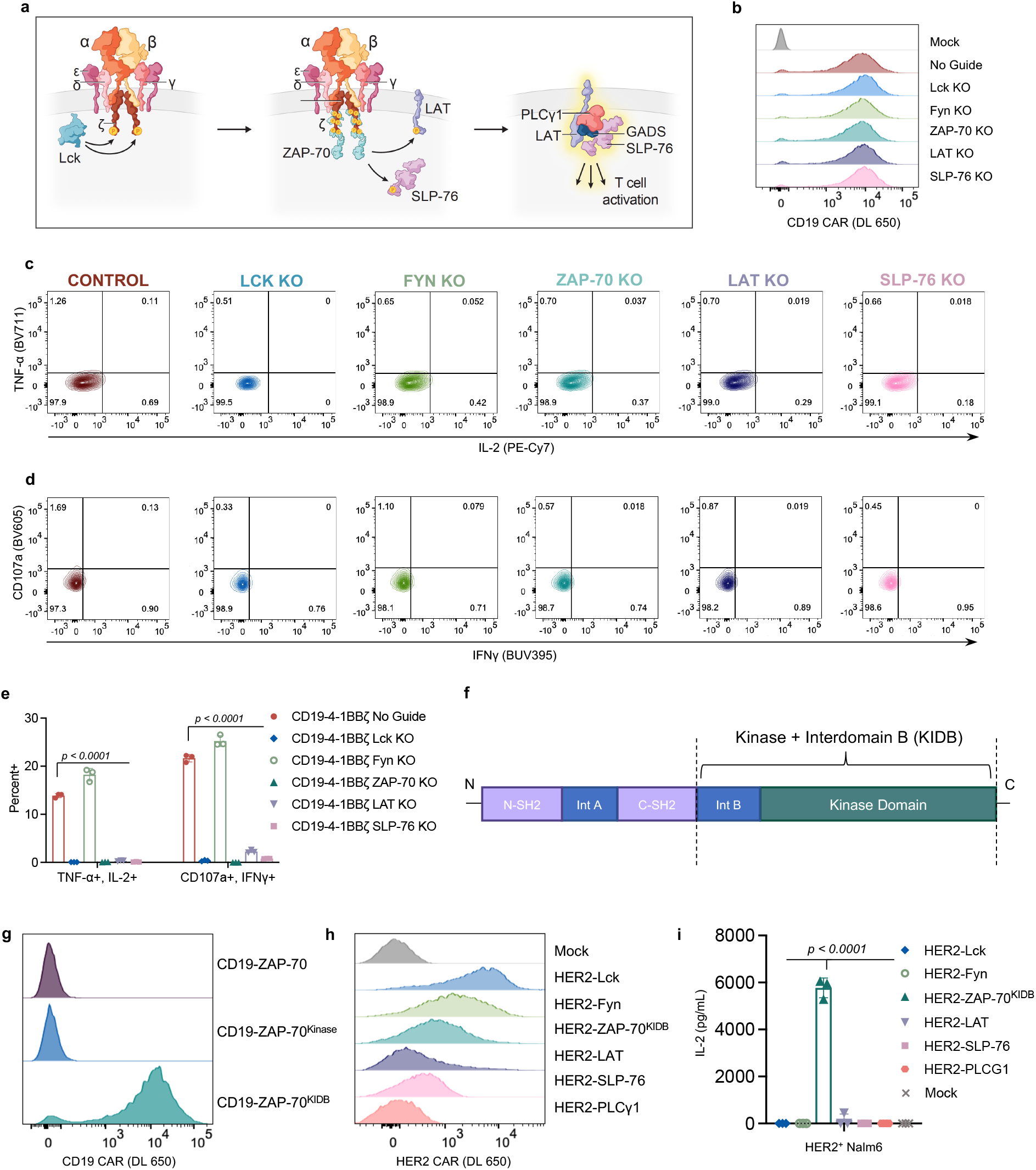
Essential Proximal Signaling Molecules for CD19-4-1BBζ CAR signal. **a**, Schematic illustrating the TCR signaling pathway wherein Lck phosphorylates ITAM motifs in CD3ζ, creating a binding site for ZAP-70. ZAP-70 is then activated and phosphorylates adapter proteins LAT and SLP-76. LAT and SLP-76 then form a scaffold for recruitment of PLCγ1 and other downstream effector molecules that initiate T cell activation. **b**, Flow cytometric data exhibiting CD19-4-1BBζ CAR expression in unedited and edited T cells prior to stimulation. **c-d**, Flow cytometric plots of TNF-α x IL-2 (**c**) and CD107a x IFNγ (**d**) in unstimulated unedited and edited CD19-4-1BBζ CAR T cells. Data is representative of three independent experiments performed with different blood donors. **e**, Quantification of TNF-α^+^IL-2^+^ and CD107a^+^IFNγ^+^ populations as shown in **Figure 1d-e**. Baseline measurements from the unstimulated controls were subtracted from stimulated conditions. Shown are mean values ± s.d. of three experimental replicates. *p* values were obtained by one way ANOVA with multiple comparisons. **f**, Schematic illustrating the protein domains of ZAP-70; dashed lines indicate the Kinase + Interdomain B (KIDB) region contained in the ZAP-70^KIDB^ CAR. **g**, Flow cytometric data exhibiting expression of CD19-ZAP-70, CD19-ZAP-70^Kinase^, and CD19-ZAP-70^KIDB^ CARs. **h**, Flow cytometric data exhibiting expression of HER2-targeting proximal signaling CARs. **i**, IL-2 secretion (as measured by ELISA) by HER2-targeting proximal signaling CARs following co-culture with HER2^+^Nalm6 tumor cells. Shown are mean values ± s.d. of three experimental replicates. Representative of three independent experiments with different T cell donors. *p* values were obtained by one way ANOVA with multiple comparisons.

**Extended Data Figure 2:**
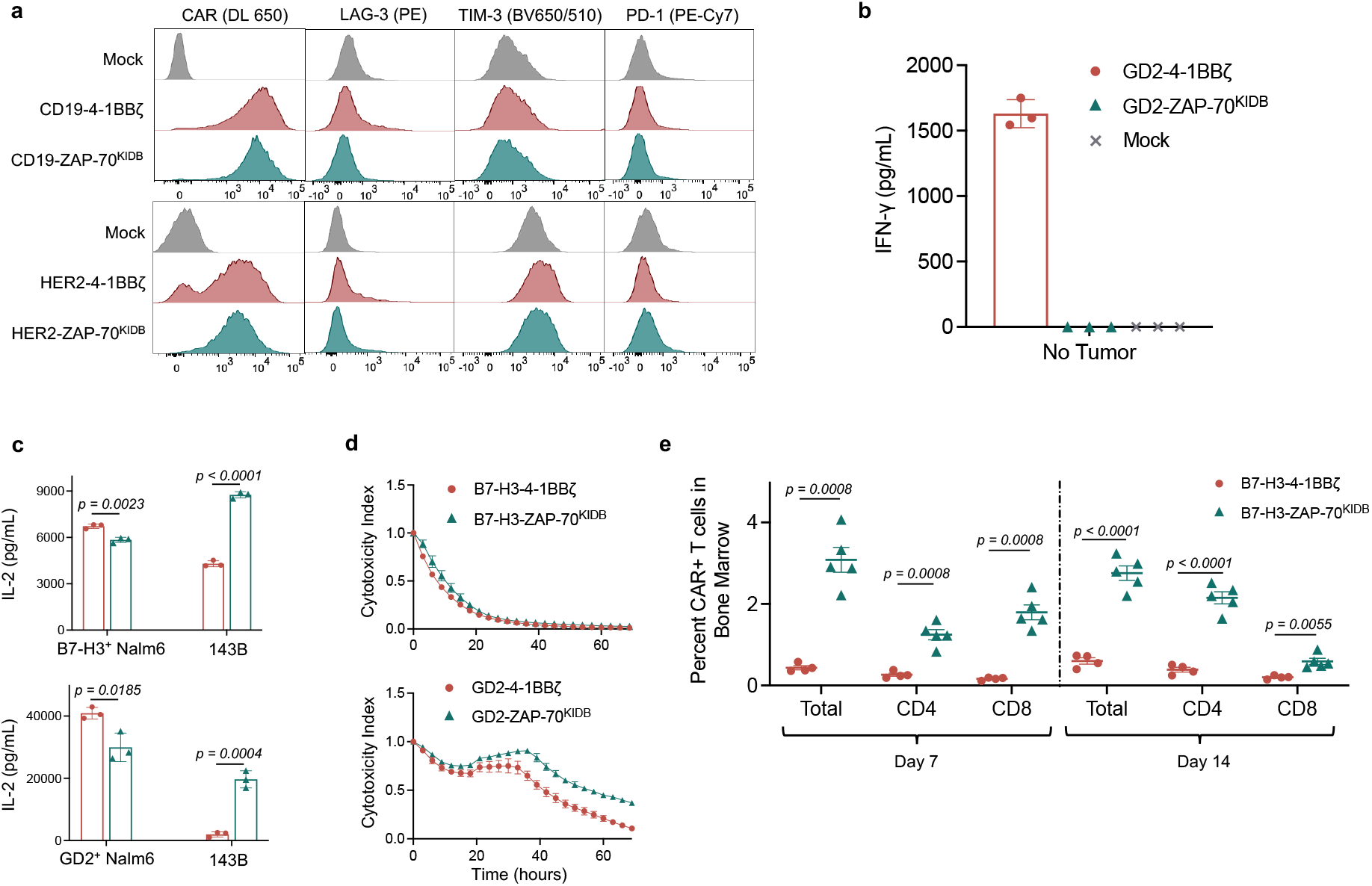
ZAP-70 CAR T cells demonstrate potent *in vitro* and *in vivo* activity. **a**, Representative flow cytometric plots of CAR, LAG-3, TIM-3, and PD-1 expression for T cells bearing CD19 or HER2-specific CARs containing 4-1BBζ or ZAP-70^KIDB^ endodomains (day 10 after T cell activation). Representative of two experiments with different T cell donors. **b**, IFNγ secretion (as measured by ELISA) by GD2-4-1BBζ and GD2-ZAP-70^KIDB^ CAR T cells following 24hr culture in the absence of target cells. Shown are mean values ± s.d. of three experimental replicates. Representative of three experiments with different T cell donors. **c**, IL-2 secretion (as measured by ELISA) by B7-H3 or GD2-specific CAR T cells containing ZAP-70^KIDB^ or 4-1BBζ endodomains following co-culture with Nalm-6 cells expressing B7-H3/GD2 or 143B osteosarcoma cells. Shown are mean values ± s.d. of three experimental replicates. Representative of four independent experiments with different T cell donors. *p* values were determined by the unpaired t-test (two-tailed). **d**, Tumor cell killing of GFP^+^ human neuroblastoma CHLA-255 cells co-cultured with B7-H3 or GD2-specific CAR T cells containing ZAP-70^KIDB^ or 4-1BBζ endodomains for at a 1:1 ratio of T cells to tumor cells. Shown are mean values ±s.d. of three experimental replicates. Representative of four independent experiments with different T cell donors. **e**, Percent CAR^+^ T cells recovered from the bone marrow of CHLA-255-bearing mice on days 7 and 14 after treatment with B7-H3-4-1BBζ or B7-H3-ZAP-70^KIDB^ CAR T cells. Shown are mean values ± s.e.m. for n=4-5 mice per group per timepoint. Experiment was performed once at two timepoints. *p* values were determined by unpaired t-test (two-tailed) with Welch’s correction.

**Extended Data Figure 3:**
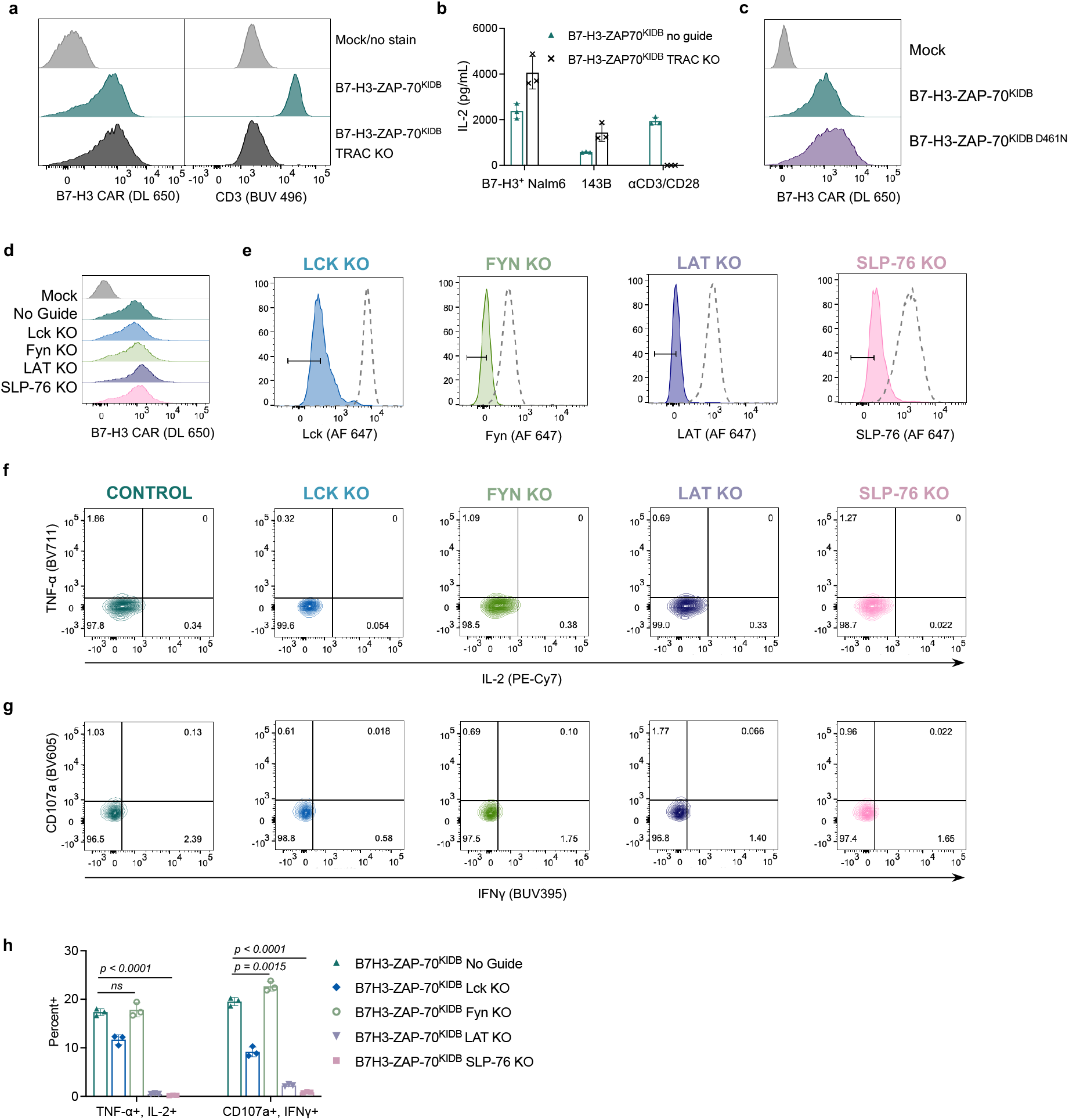
Mechanisms of signaling in B7-H3-ZAP-70^KIDB^ CAR T cells. **a**, Flow cytometric data exhibiting surface CAR and CD3 expression of B7-H3-ZAP-70 (± TRAC knockout) **b**, IL-2 secretion (as measured by ELISA) by B7-H3-ZAP-70 (± TRAC knockout) CAR T cells shown in **a**. Shown are mean values ± s.d. of three experimental replicates. Representative of two experiments with different T cell donors. **c**, Flow cytometric data exhibiting the expression of B7-H3-ZAP-70^KIDB^ CARs ± kinase-ablating D461N mutation. **d**, Flow cytometric data exhibiting B7-H3-ZAP-70^KIDB^ CAR expression in unedited and edited T cells prior to stimulation. **e**, Flow cytometric plots demonstrating knockout efficiencies for proximal signaling molecules in CAR T cells shown in **d**. **f-g**, Flow cytometric plots of TNF-α x IL-2 (**f**) and CD107a x IFNγ (**g**) in unstimulated unedited and edited B7-H3-ZAP-70^KIDB^ CAR T cells. Data is representative of three independent experiments performed with two different T cell donors. **h,** Quantification of TNF-α^+^IL-2^+^ and CD107a^+^IFNγ^+^ populations as shown in **Figure 2h-i**. Baseline measurements from the unstimulated controls were subtracted from the stimulated conditions. Shown are mean values ± s.d. of three experimental replicates. *p* values were obtained by one way ANOVA with multiple comparisons.

**Extended Data Figure 4:**
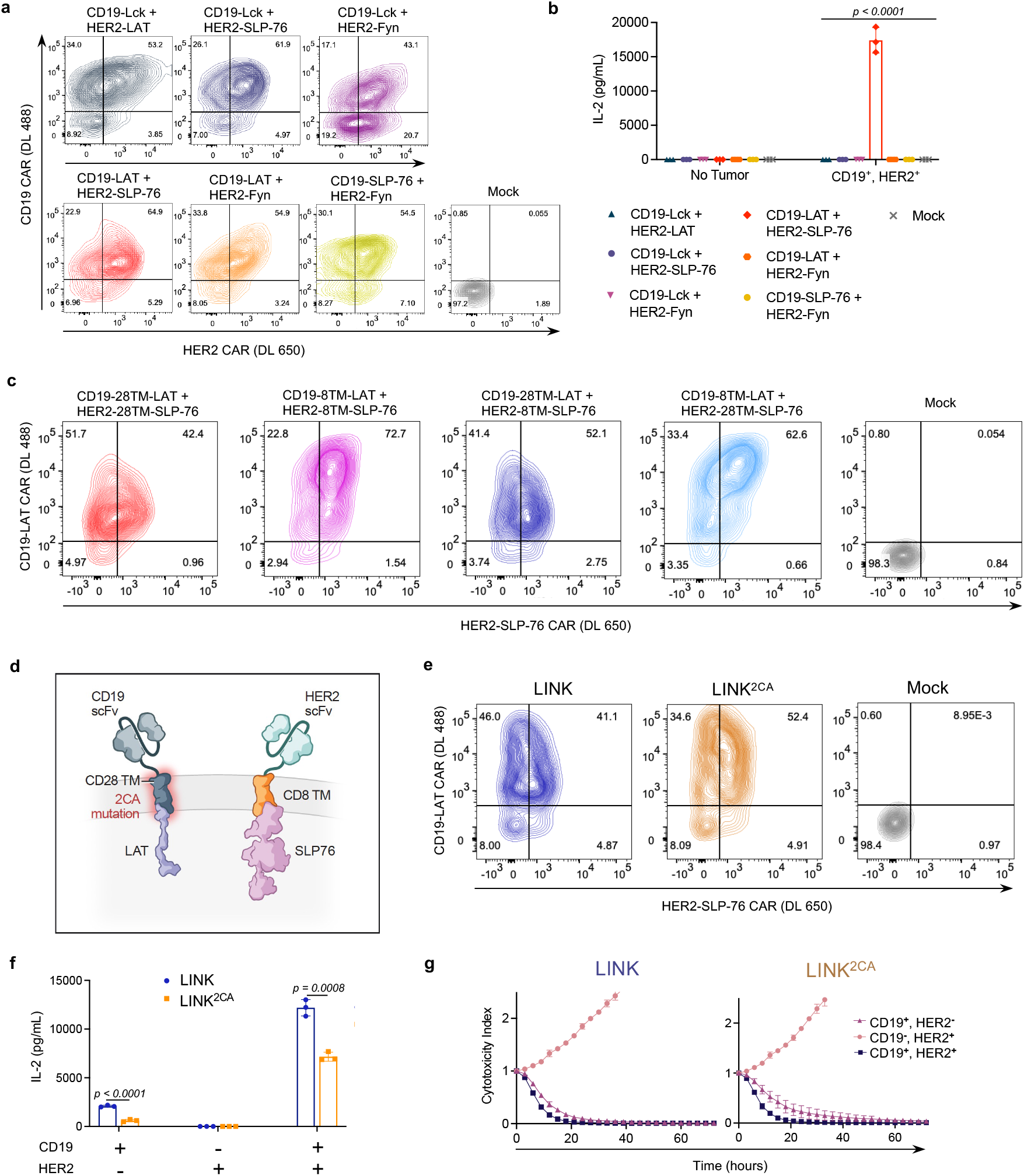
LAT and SLP-76 CARs jointly mediate T cell activation. **a**, Flow cytometric data exhibiting CAR expression of co-transduced CD19 and Her2 proximal signaling CAR (Lck, Fyn, LAT, and SLP-76) combinations. **b,** IL-2 secretion (as measured by ELISA) by T cells from **a** following coculture with HER2^+^Nalm6 (CD19^+^, HER2^+^) cells. Shown are mean values ± s.d. of three experimental replicates. Representative of four experiments performed with three different T cell donors. *p* values were obtained by one way ANOVA with multiple comparisons. **c**, Flow cytometric expression of LAT and SLP-76 CARs on T cells utilized in assays in **Figure 3f-g** on. **d**, Schematic illustrating incorporation of a dual Cysteine-to-Alanine (2CA) mutation in the CD28 hinge-transmembrane (TM) domain of the LAT CAR component of the LINK system. **e**, Flow cytometric expression of LINK CAR components (±2CA mutation). **f,** IL-2 secretion (as measured by ELISA) by LINK CAR T cells (±2CA mutation) following coculture with cell lines shown in **Figure 3c**. Shown are mean values ± s.d. of three experimental replicates. Representative of eight independent experiments performed with five different T cell donors. *p* values were obtained by unpaired two-tailed t-tests. **g,** Tumor cell killing of cell lines shown in **Figure 3c** co-cultured with LINK CAR T cells (±2CA mutation) at a 2:1 ratio of T cells to tumor cells. Shown are mean values ± s.d. of three experimental replicates. Representative of eight independent experiments performed with five different T cell donors.

**Extended Data Figure 5:**
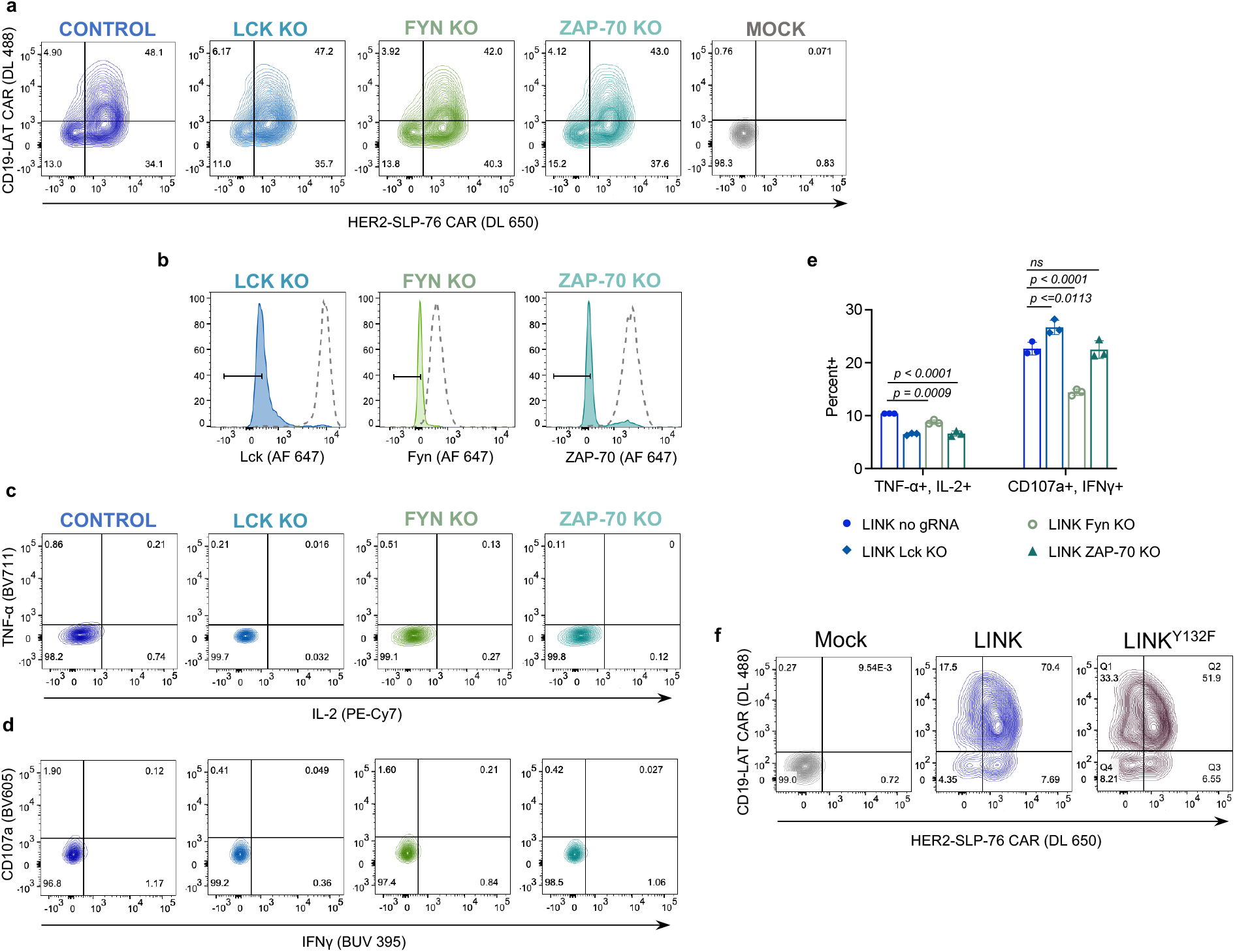
Knockout of proximal signaling proteins in LINK CAR T cells. **a**, Flow cytometric data exhibiting LINK CAR (CD19-28TM-LAT + HER2-8TM-SLP-76) expression in unedited and edited T cells prior to stimulation with HER2^+^ Nalm6. **b,** Flow cytometric plots demonstrating knockout efficiencies for proximal signaling molecules in CAR T cells shown in **a**. **c-d,** Flow cytometric plots of TNF-α x IL-2 (**c**) and CD107a x IFNγ (**d**) in unstimulated unedited and edited LINK CAR T cells. Data is representative of two independent experiments performed with different T cell donors. **e,** Quantification of TNF-α^+^IL-2^+^ and CD107a^+^IFNγ^+^ populations as shown in **Figure 3h-i**. Baseline measurements from the unstimulated controls were subtracted from the stimulated conditions. Shown are mean values ± s.d. of three experimental replicates. *p* values were obtained by one way ANOVA with multiple comparisons. **f,** Flow cytometric expression of LINK CAR (±LAT ^Y132F^).

**Extended Data Figure 6:**
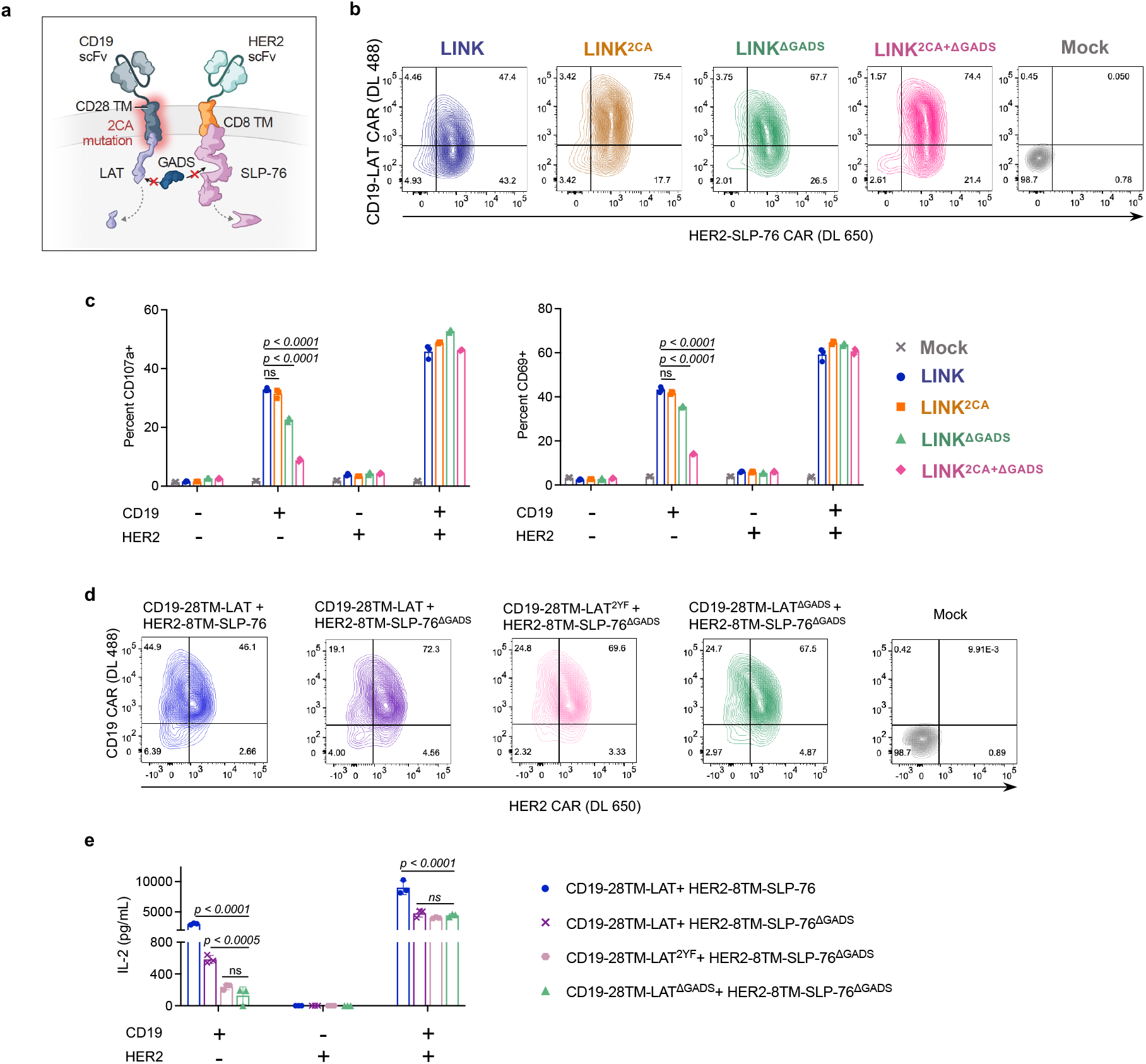
Disrupting GADS interactions eliminates LINK CAR leakiness. **a**, Schematic illustrating LINK CAR bearing both Cysteine-to-Alanine (2CA) mutations and GADS binding site deletions (ΔGADS). **b**, Flow cytometric expression of LINK CARs (±2CA, ±ΔGADS) on T cells utilized in assays in **Figure 4b-d**. **c**, Quantification of CD 107a^+^ and CD69 on indicated LINK CAR T cells following co-culture with cell lines shown in **Figure 3c**. Baseline measurements from the CD107a unstimulated controls were subtracted from the stimulated CD107a conditions. Representative of five independent experiments with four different T cell donors. Shown are mean values ± s.d. of three experimental replicates. *p* values were obtained by one way ANOVA with multiple comparisons. **d**, Flow cytometric expression of LINK CARs bearing either Y171F/Y191F point mutations (2YF) or truncation of the GADS binding regions (ΔGADS). **e**, IL-2 secretion (as measured by ELISA) by indicated LINK CAR T cells following co-culture with cell lines shown in **Figure 3c**. Shown are mean values ± s.d. of three experimental replicates. Performed one time. Note that data for CD19-28TM-LAT + HER2-8TM-SLP-76 and CD19-28TM-LAT^ΔGADS^+ HER2-8TM-SLP-76^ΔGADS^ conditions are identical to **Figure 4b**. *p* values were obtained by one way ANOVA with multiple comparisons.

**Extended Data Figure 7:**
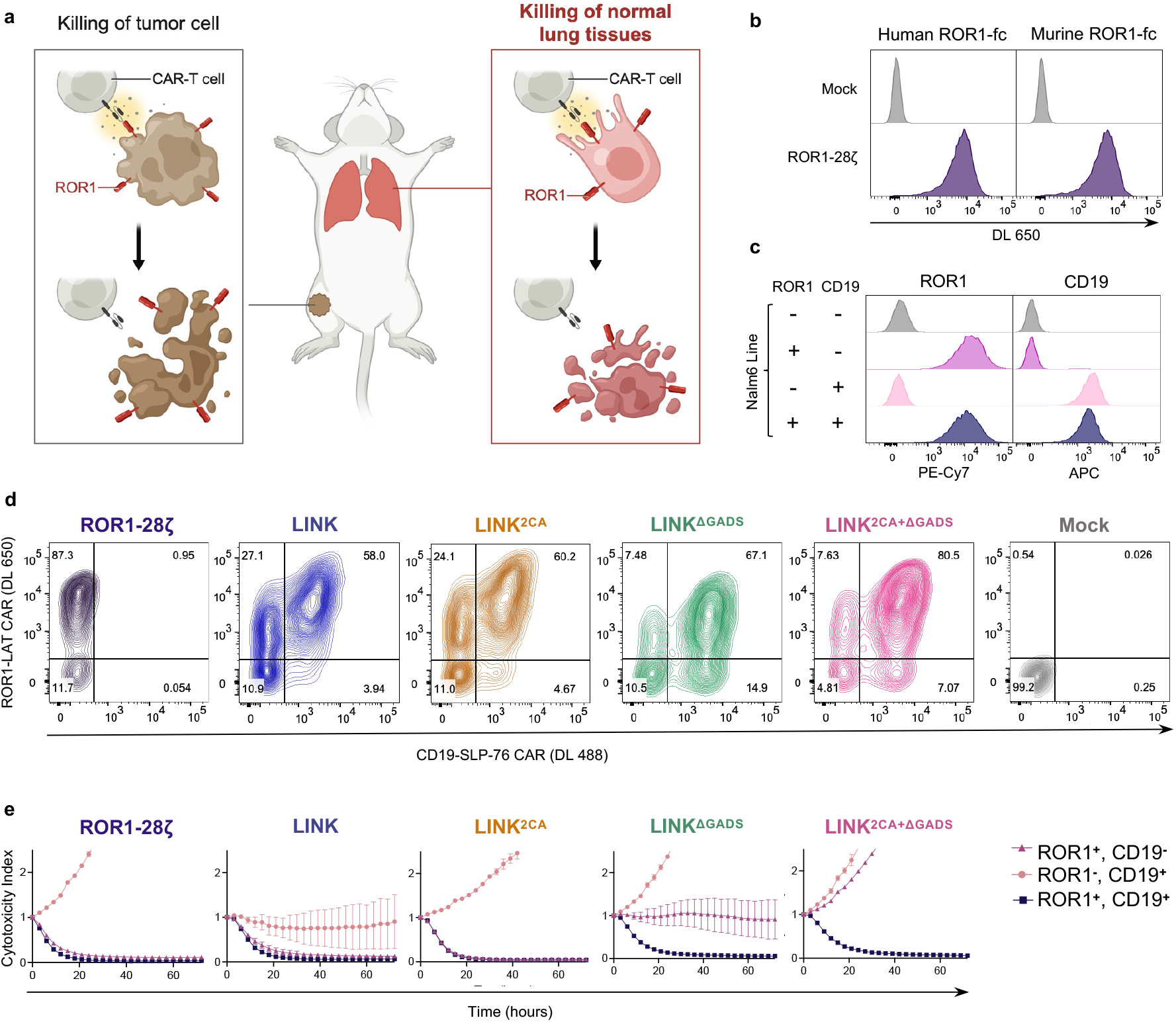
Design of a ROR1-targeting LINK CAR for testing in a model of on-target, off-tumor toxicity. **a**, Schematic illustrating on-target, off-tumor toxicity in the lungs of tumor-bearing mice treated with ROR1 targeted CAR T cells. **b**, Flow cytometry plots exhibiting detection of ROR1-28ζ CAR on T cells with both recombinant human and murine ROR1. **c**, ROR1 and CD19 expression on single/double antigen positive Nalm6 lines used for AND-gate verification experiments. **d**, Flow cytometric expression of ROR1-28ζ or indicated ROR1/CD19-targeted LINK CARs on T cells used for *in vitro* and *in vivo*(**Figure 4e-g**) testing. **e**, Tumor cell killing of cell lines shown in **c** co-cultured with the indicated CAR T cells at a 2:1 ratio of T cells to tumor cells. Shown are mean values ± s.d. of three experimental replicates. Representative of four independent experiments performed with two different T cell donors.

**Extended Data Figure 8:**
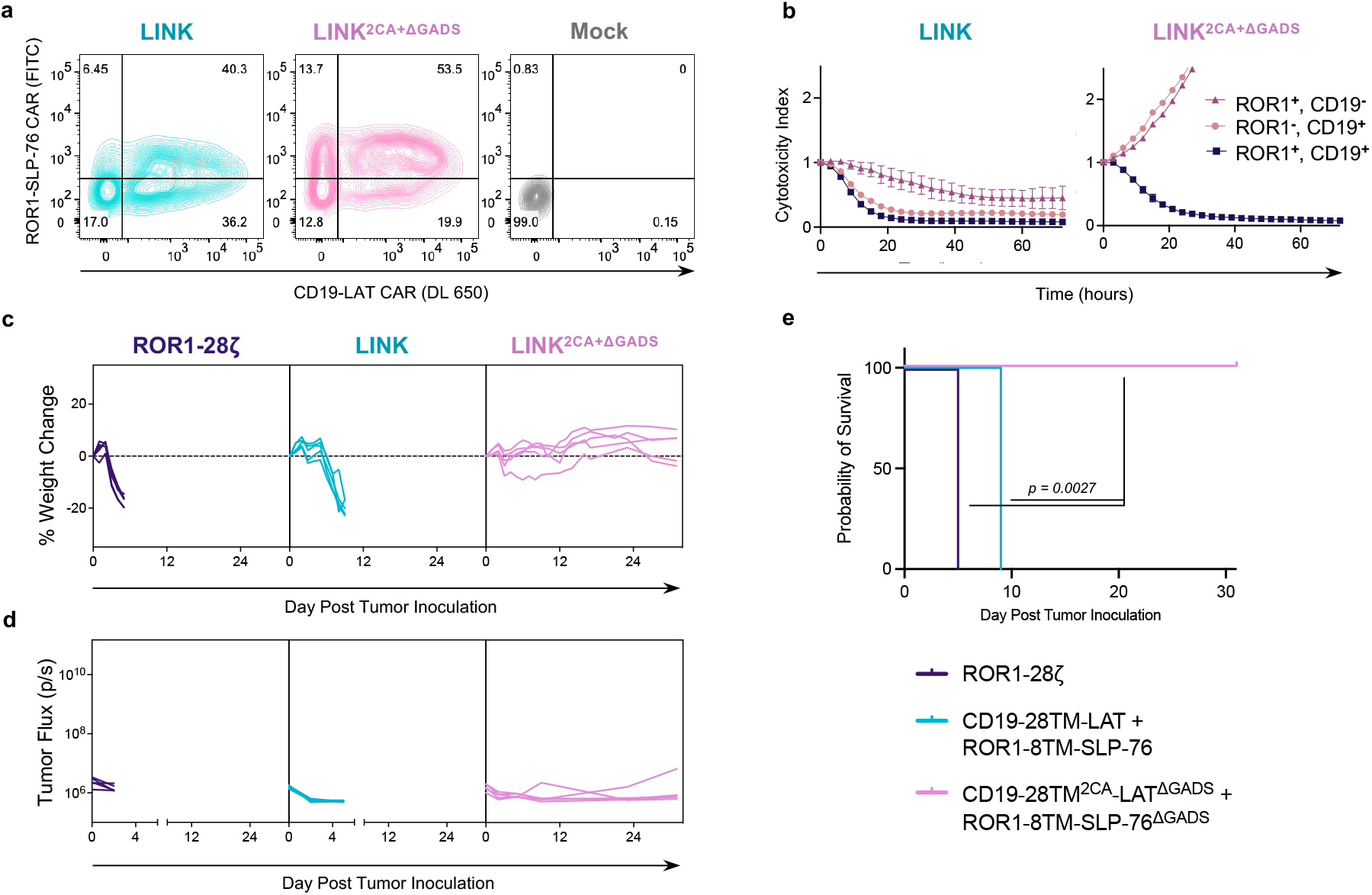
Elimination of single antigen reactivity is essential for both LINK CAR components. **a,** Flow cytometric expression of ROR1/CD19-targeted LINK CARs on T cells. **b**, Tumor cell killing of indicated cell lines by the LINK CAR T cells at a 2:1 ratio of T cells to tumor cells. Shown are mean values ± s.d. of three experimental replicates. Performed one time. **c-e**, NSG mice bearing ROR1^+^Nalm6-luciferase were treated with the indicated LINK CAR T cells with the SLP-76 CAR bearing specificity for ROR1 (reversed from Figure 4). **(c)** Weights for individual mice over time plotted as a percentage of the weight on day 0. **(d)** Quantification of tumor progression for each individual mouse as measured by flux values acquired via bioluminescence imaging (BLI). (**e**) Survival curves for mice bearing tumors shown in **d.** *p* values were determined by the Log-rank test. Performed once with n=5 mice per group.

**Extended Data Figure 9:**
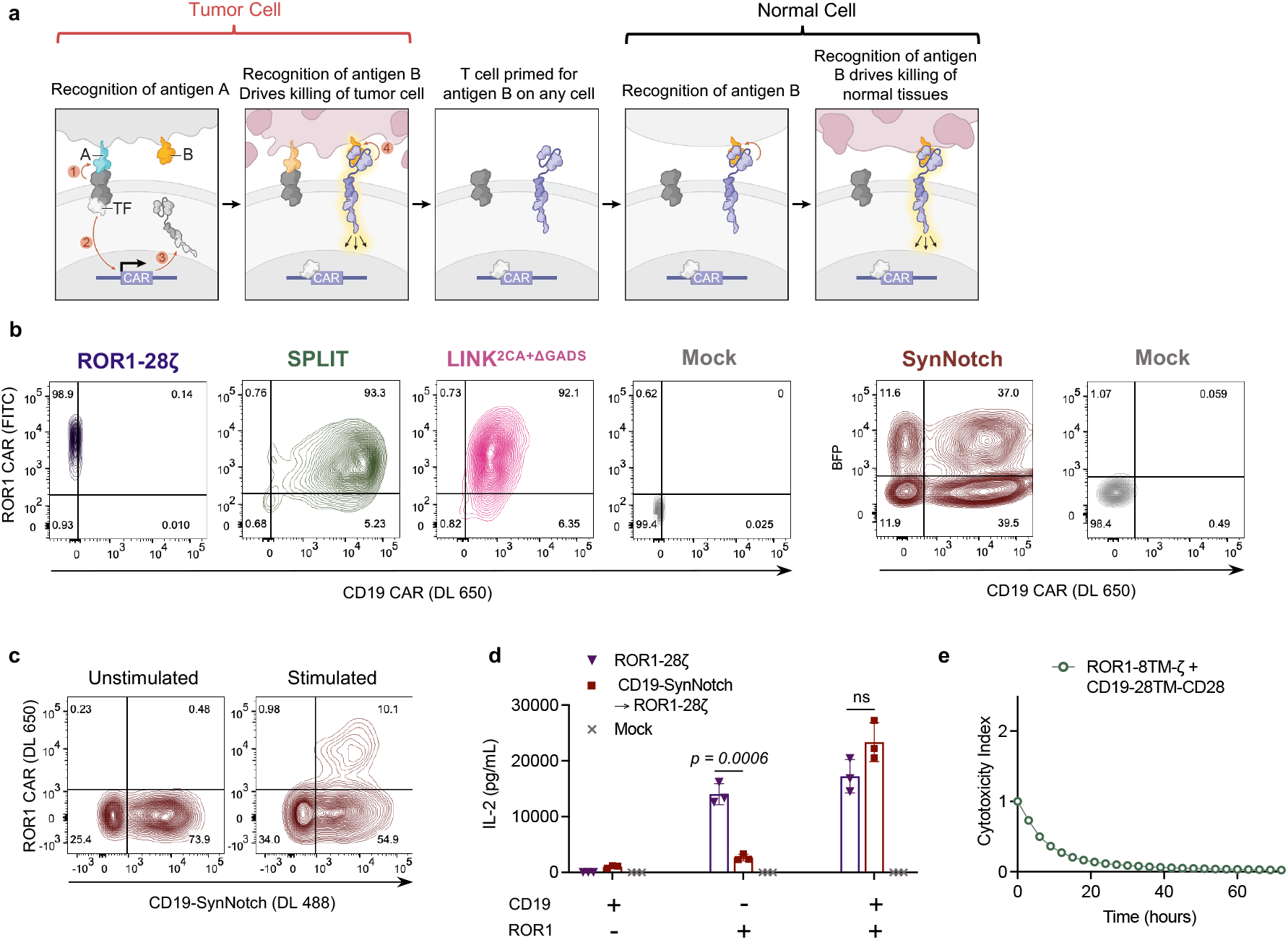
LINK CAR outperforms both SynNotch and SPLIT CAR systems. **a**, Schematic illustrating the potential for on-target, off-tumor toxicity for SynNotch CAR T cells. **b**, Flow cytometric expression of ROR1-28ζ and ROR1/CD19-targeted LINK, SPLIT, and SynNotch on T cells day 10 after activation. **c**, Inducible ROR1-28ζ CAR expression (detected with recombinant ROR1 protein) following stimulation of the CD19-SynNotch receptor through 24 hour stimulation with Nalm6. **d**, IL-2 secretion (as measured by ELISA) by ROR1-28ζ and CD19-SynNotch → ROR1-28ζ CAR T cells following co-culture with the indicated cell lines. Shown are mean values ± s.d. of three experimental replicates. Conducted with the same T cells used *in vivo* for **Figure 4h-j**. *p* values were obtained by unpaired two-tailed t-tests. **e**, Killing of ROR1^+^Nalm6 (CD19^+^, ROR1^+^) tumor cells by ROR1/CD19-targeting SPLIT CAR T cells cocultured at a 2:1 T cell to tumor ratio. Shown are mean values ± s.d. of three experimental replicates. Representative of two independent experiments with different T cell donors.

**Extended Data Figure 10:**
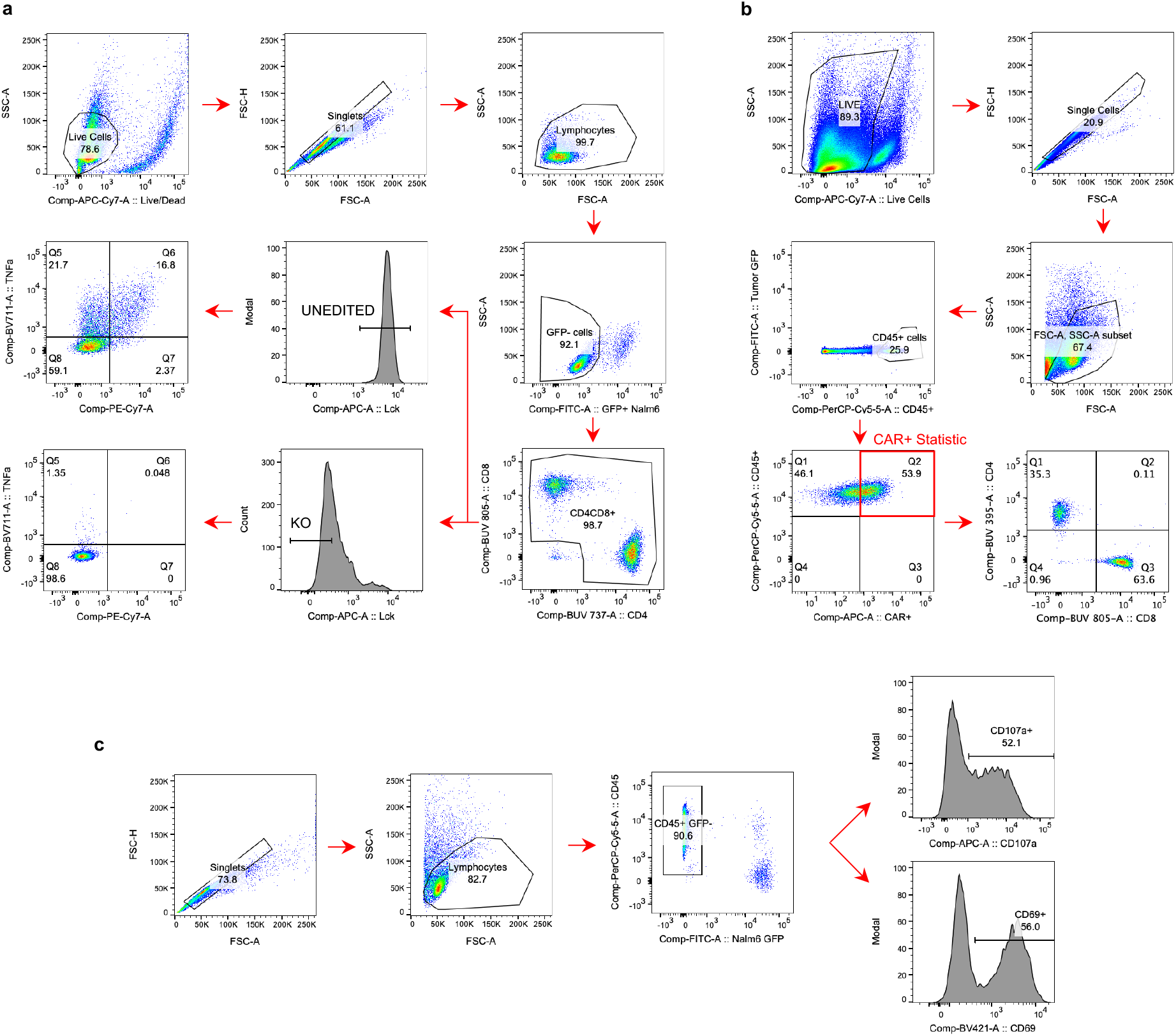
Gating strategies for flow cytometry. **a,** Intracellular cytokine staining of CRISPR/Cas9-edited CAR T Cells (Figures **1c-e**, **2h-i**, **3h-i**; Extended Data Figures **1c-e**, **3e-h**, **5b-e**). **b,** Detection of CAR^+^ T cells isolated from murine spleen and bone marrow tissue (Figure **2e**; Extended Data Figure **2e**). **c,** LINK CAR T cell activation assays measuring CD107a and CD69 (Figure **4d**; Extended Data Figure **6c**).

